# CpGPT: a Foundation Model for DNA Methylation

**DOI:** 10.1101/2024.10.24.619766

**Authors:** Lucas Paulo de Lima Camillo, Raghav Sehgal, Jenel Armstrong, Max Melnikas, Henry E. Miller, Jun Ding, Luigi Ferruci, Jessica A. Lasky-Su, Albert T. Higgins-Chen, Steve Horvath, Bo Wang

## Abstract

DNA methylation is a type of epigenetic modification that plays a significant role in development, aging, and disease. Despite extensive research, how genome-wide DNA methylation patterns collectively encode and influence complex phenotypes such as aging and disease remains difficult to characterize with conventional approaches. Foundation models are a class of machine learning model that leverage vast quantities of data to make sense of complex data types, such as genome sequences or single-cell transcriptomes. Here, we present the Cytosine-phosphate-Guanine Pretrained Transformer (CpGPT), a novel foundation model pretrained on CpGCorpus, a novel database with more than 2,000 DNA methylation datasets encompassing over 150,000 samples from diverse conditions. CpGPT leverages an improved transformer architecture to learn comprehensive representations of methylation patterns, allowing it to impute and reconstruct genome-wide methylation profiles from limited input data. By capturing sequence, positional, and epigenetic contexts, CpGPT outperforms specialized models when finetuned for aging-related tasks, including the state-of-the-art GrimAge2 and PCGrimAge for mortality and morbidity estimation. The model is highly adaptable and can impute beta values across different methylation platforms, tissue types, mammalian species, and even single-cell data. As a foundation model, CpGPT can be leveraged as a new tool for biological discovery in the field of epigenetics. The open-source code and model can be found at http://github.com/lucascamillomd/CpGPT.

**Highlights:**

- CpGPT is a novel foundation model for DNA methylation analysis, pretrained on over 2,000 datasets encompassing 150,000+ samples.
- The model demonstrates strong performance in zero-shot tasks including imputation, array conversion, and reference mapping.
- CpGPT achieves state-of-the-art results in mortality prediction and chronological age estimation.

## 1 Introduction

Since the introduction of the transformer architecture [1], artificial intelligence has undergone rapid advancements, particularly due to the development of foundation models and large language models (LLMs) [2]. Transformers leverage self-attention mechanisms to process sequential data more effectively, capturing long-range dependencies and complex patterns [3]. Pretrained on vast amounts of data in an unsupervised manner, these models have demonstrated exceptional performance across a variety of downstream tasks through transfer learning, making them highly versatile and effective.

Beyond natural language processing, transformers and foundation models have significantly impacted biology and medicine. They have advanced the analysis of single-cell transcriptomic data, uncovering previously unknown biology. Models such as scGPT [4], Geneformer [5], and Universal Cell Embeddings [6] display state-of-the-art performance for several tasks and even possess emergent behavior. The ability of transformers to integrate sequence, structural, and contextual information makes them particularly suitable for biological data, which often involve complex interactions and hierarchies. The advent of these models has also begun to influence longevity research [7, 8].

Despite significant progress in the past decade in the field of epigenetic biomarkers, many widely used models rely on relatively simple regularized linear models using Cytosine-phosphate-Guanine (CpG) DNA methylation data [9, 10, 11, 12, 13]. These predictors often do not consider the sequence context or genomic positions of CpG sites and as a result, they may overlook complex interactions and the underlying biological mechanisms driving processes like development, aging, and disease. A recent advancement involves the application of principal component decomposition before the linear model to enhance the reliability and performance of DNA methylation epigenetic proxies [14, 15]. However, few predictors, such as AltumAge [16] and DeepMAge [17], have utilized deep neural networks to model the complex relationships within methylation data.

Motivated by these advances, we have developed the CpG Pretrained Transformer (CpGPT), a transformer-based deep neural network that leverages the attention mechanism to effectively learn relationships between methylation sites by incorporating sequence, positional, and epigenetic information. As a foundation model, CpGPT is capable of performing a series of tasks in both zero-shot settings and when finetuned. The model can impute missing methylation values within a dataset, convert between different methylation platforms by reconstructing unmeasured CpG sites, and perform zero-shot reference mapping to label samples without finetuning, all while benefiting from increasing test-time compute. Moreover, CpGPT excels when finetuned for specific tasks, achieving second place overall for chronological age prediction in the Biomarkers of Aging Challenge [18], and outperforming the state-of-the-art GrimAge2 for mortality estimation across independent cohorts. Concurrent with our work, MethylGPT was developed as an independent foundation model for DNA methylation, further underscoring the growing interest in transformer-based approaches for epigenetic analysis [19].

The CpGPT framework is highly generalizable and can be applied across various tasks, such as cancer prediction and classification, across different mammalian species, and across platforms, including single-cell data. CpGPT establishes a new standard for DNA methylation analysis and offers a versatile tool for multiple applications in the field of epigenetics.

## 2 Results

### 2.1 Developing a foundation model for DNA methylation

To fully harness the capabilities of foundation models, which often improve with increased data availability, we first curated a large-scale database named *CpGCorpus* (see Methods). This database comprises over 150,000 human DNA methylation samples collected from more than 2,000 studies, encompassing a diverse array of tissue types, developmental stages, and disease conditions (Supplementary Figure 1). We preprocessed and harmonized the data to ensure consistency across different methylation array platforms, including Illumina 27k, 450k, EPIC, and EPICv2 arrays, resulting in over 101 billion tokens, where each token is one CpG measurement in one sample.

Building upon this extensive dataset, we designed the CpGPT model architecture to capture three primary types of contextual information: (1) sequence-based context; (2) local and global positional context; and (3) epigenetic state. To encode the sequence context, we utilized embeddings of the 1000 flanking nucleotides around each CpG site (2001 base pairs in total) derived from a pretrained DNA language model, namely Nucleotide Transformer V2 with 500 million parameters (NTv2_500M_) [20]. For global positional context, we sorted the sequence embeddings by genomic positions and grouped them by chromosomes. As each input is a variable-size subset of all CpG sites, we applied stochastic shuffling of the chromosome order during training to prevent the model from associating a fixed input position with a specific CpG locus and learning spurious positional shortcuts. We incorporated modifications of the original positional encoding method from Vaswani et al. [1] along with rotary positional embeddings [21] to enable the model to understand both the overall genomic structure and the local relationships between methylation sites. Finally, we transformed the single-value methylation state (beta value) of each CpG site into an embedding representing its epigenetic status using a dedicated encoder. These embeddings were integrated to form the input to the model (Figure 1).

**Figure 1:**
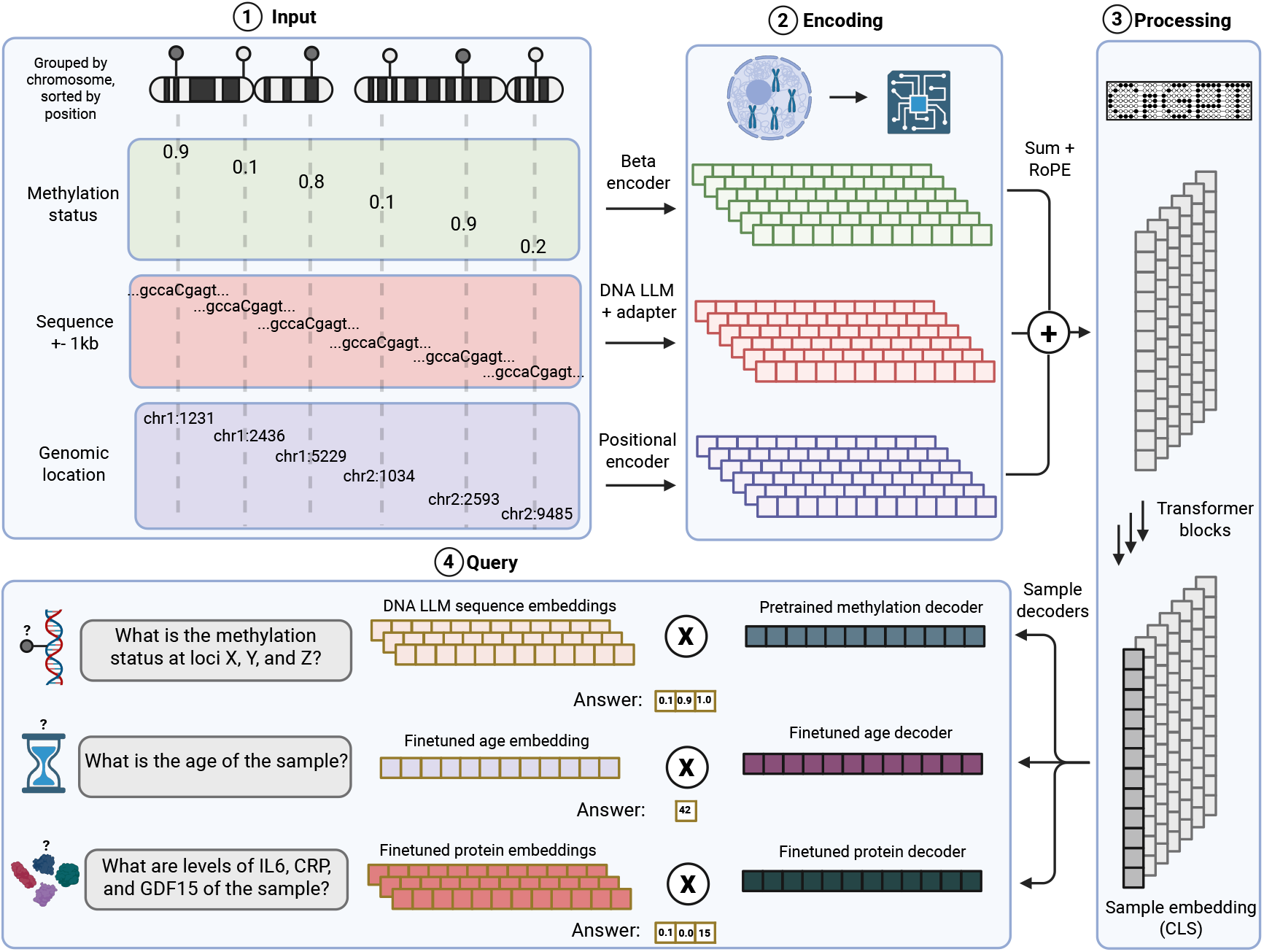
The CpGPT framework. The input CpG sites are grouped by chromosome and sorted. Information is encoded by the model through several neural networks, including a projection of the methylation status, a transformation of the sequence embedding from a DNA large language model (LLM), and a positional addition to incorporate genomic location. The embeddings of each information type are added followed by rotational positional encoding (RoPE). A series of transformer blocks aggregates the sample information into a sample embedding (CLS). The sample embedding can be queried after transformation with a pretrained or finetuned decoder followed by a dot product of the query embedding.

At the core of CpGPT is an enhanced version of the transformer architecture, known as transformer++ [22], which includes several modifications to improve model expressivity and training stability. CpGPT learns to create meaningful sample-level embeddings that capture the comprehensive methylation profile of each sample. This is achieved by training the model in an unsupervised manner to predict beta values and their associated uncertainty for CpG sites. The model can utilize the DNA LLM embedding of any genomic position, including those not seen during training, as a query to reconstruct the methylation state of that specific locus.

The training process for CpGPT involves a multitask learning approach, employing various loss functions to optimize different aspects of the model’s performance. These include losses for accurate beta value prediction, uncertainty estimation, and the quality of sample embeddings (see Methods). The model is designed to handle missing data, enabling it to work effectively with incomplete methylation profiles, which are common because of varying array designs and experimental conditions. Rather than relying on a fixed vocabulary of CpG sites as current epigenetic clocks, the model’s input is typically a random subset of CpG sites, varying from 5,000 to 10,000 depending on the task. This comprehensive training strategy allows CpGPT to learn rich, nuanced representations of DNA methylation patterns across diverse genomic contexts and biological conditions, allowing it to perform a series of downstream tasks.

We pretrained two models, a lightweight one designed to facilitate finetuning, and a large one, tailored toward zero-shot tasks. The former contains 2.5 million parameters, whereas the latter contains 101 million. They are herein referred to as CpGPT_2M_ and CpGPT_100M_ respectively. Throughout this paper, we adopt a systematic naming convention for finetuned variants: CpGPT_size_-Task, where the subscript denotes the pretrained model size and the suffix after the hyphen indicates the downstream task.

To validate these architectural decisions, we conducted systematic ablations varying the DNA LLM backbone, context window length, transformer architecture, input ordering strategy, and training duration (Supplementary Figure 2; Supplementary Tables 1–3). We compared three sizes of two architectures of DNA sequence encoders (HyenaDNA, NTv2) across three flanking context lengths, all trained for 100,000 steps [23, 20]. The largest NTv2_500M_ with a 2001 bp context yielded the lowest test beta MAE of 0.0704, outperforming the biggest HyenaDNA_large_ variant with MAE = 0.0794. Longer contexts consistently improved performance across all backbones, with diminishing returns between 1001 and 2001 bp. Removing the transformer blocks while keeping all other components increased the test beta MAE from 0.0704 to 0.0802 (+14.0%), demonstrating that the self-attention mechanism provides a meaningful contribution even at the 2.5M parameter scale. Replacing chromosomal positional sorting with random CpG ordering resulted in marginally worse performance (MAE = 0.0709). Finally, we tracked performance from 20k to 1M gradient steps and observed continued improvement following an approximate power-law scaling curve, with the test beta MAE decreasing from 0.0931 at 20,000 steps to 0.0613 at one million steps.

### 2.2 CpGPT learns meaningful methylation and sample embeddings

Understanding how well a foundation model captures the underlying structure of biological data is essential. We assessed whether CpGPT can learn meaningful embeddings in an unsupervised manner by examining whether 1. features (individual CpG sites) and 2. samples naturally cluster according to their biological attributes, even without explicit labels. To this end, we generated low-dimensional representations of the high-dimensional embeddings using Uniform Manifold Approximation and Projection (UMAP) [24] at both the CpG site (locus) level and the sample level across a variety of datasets.

Our first experiment aimed to test whether the locus embeddings produced by CpGPT capture biologically relevant genomic annotations. Specifically, CpGPT transforms the NTv2_500M_ embeddings of nucleotide sequences flanking each CpG site using an adapter network to match CpGPT’s internal dimension. We hypothesized that these transformed locus embeddings would better reflect functional genomic annotations than the raw DNA LLM embeddings, given the model’s knowledge of epigenetics. To test this, we sampled approximately 100,000 CpG sites from the Illumina EPICv2 array and applied UMAP for visualization. We then examined the resulting clusters with chromatin state annotations obtained from the KnowYourCG (KYCG) EPICv2 knowledgebase [25], which provides consensus ChromHMM [26] segmentations derived from ENCODE data. These consensus annotations integrate chromatin state calls across multiple cell types and tissues, yielding a tissue-agnostic reference that segments the genome into distinct states such as active transcription start sites (TssA), bivalent promoters (TssBiv; characterized by both activating and repressive marks often seen in developmentally regulated genes), enhancers (Enh), and repressed or inactive regions. In the UMAP projection of the CpGPT_100M_ locus embeddings (Figure 2a), CpG sites linked to active TssA regions formed a distinct cluster, while bivalent promoters and enhancers occupied intermediate positions between active and repressed states. These results indicate that CpGPT’s locus embeddings segregate CpG sites according to their chromatin state, demonstrating the model’s capacity to capture complex epigenetic information solely from sequence context, particularly when compared to NTv2_500M_ (Supplementary Figure 3).

**Figure 2:**
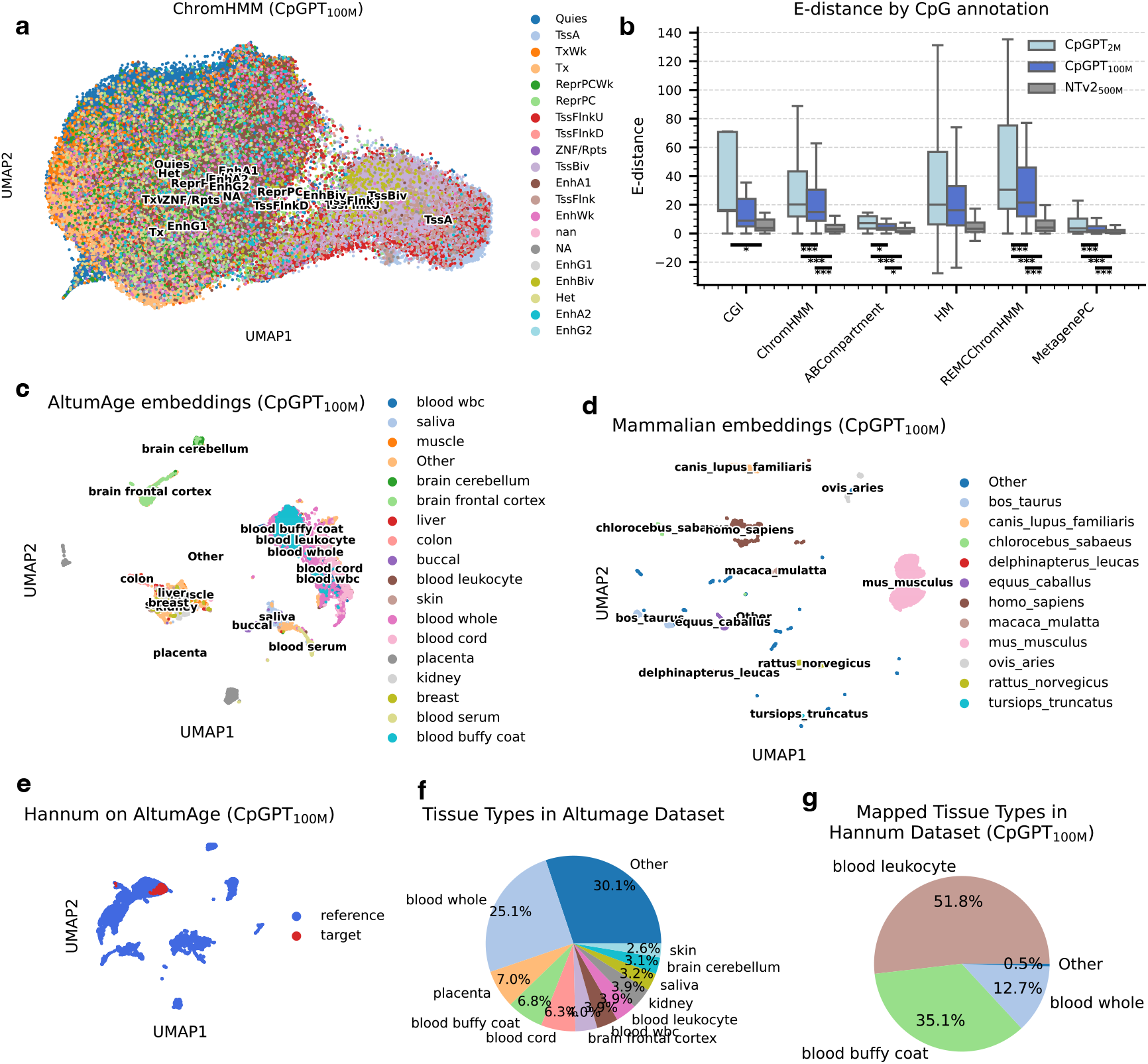
CpGPT embeddings capture biologically meaningful information. (a) UMAP of CpGPT_100M_ locus embeddings for 100,000 randomly sampled CpG sites (Illumina EPICv2 array), colored by ChromHMM annotation. (b) Boxplot showing energy distances for six genomic annotations, comparing the genomic location embeddings of CpGPT_2M_, CpGPT_100M_, and the Nucleotide Transformer V2 500M. Mann-Whitney two-sided test p-value shown [* (p < 0.05), ** (p < 0.005), *** (p < 0.0005)]. Boxes show the first and third quartiles (IQR), whiskers extend to the last data point within 1.5 IQR, and the middle line marks the median. (c) UMAP of CpGPT_100M_ sample embeddings for the AltumAge dataset, colored by tissue type (with “other” indicating tissue labels totaling less than 10% of the data). (d) UMAP of CpGPT_100M_ sample embeddings for 55 mammalian species (GSE223748), colored by species (with “other” indicating species labels totaling less than 10% of the data). (e) UMAP of CpGPT_100M_ sample embeddings comparing the AltumAge dataset (reference in blue) and the Hannum dataset (target in red). (f) Pie chart illustrating the tissue composition of the AltumAge dataset. (G) Pie chart illustrating the tissue composition of the Hannum dataset.

In order to quantitatively validate our visual observations, we analysed six genomic annotation frameworks (CGI, ChromHMM, ABCompartment, Histone Modifications, REMC-ChromHMM [27], and MetagenePC) for each embedding type across three complementary metrics: energy distance (E-distance) [28], Adjusted Rand Index (ARI), and Normalized Mutual Information (NMI) (see Methods). E-distance measures pairwise group separation, while ARI and NMI compare KMeans-discovered clusters against ground-truth annotation labels. CpGPT_2M_ achieved the best average rank (1.22 out of 3), followed by CpGPT_100M_ (2.28) and NTv2_500M_ (2.50) (Supplementary Table 4). The advantage was particularly pronounced for chromatin-state annotations: for ChromHMM, CpGPT_2M_ ranked first on all three metrics (ARI = 0.083, NMI = 0.237, Median E-distance = 20.28), while NTv2_500M_ ranked last (ARI = 0.054, NMI = 0.166, Median E-distance = 2.44). The E-distance results were consistent, with significantly higher values for both CpGPT variants compared to NTv2_500M_ across ChromHMM, REMCChromHMM, and MetagenePC annotations (Mann-Whitney U test, all *p* ≤ 0.0005). These findings confirm that CpGPT produces more distinctly clustered locus embeddings than the underlying DNA LLM alone.

Next, we evaluated whether the sample-level embeddings learned by CpGPT encapsulate meaningful biological variation. For this purpose, we extracted sample embeddings from several publicly available datasets: (i) a multi-tissue DNA methylation dataset previously used to develop the AltumAge clock [16], (ii) a Reduced Representation Bisulfite Sequencing (RRBS) atlas (GSE233417) [29] encompassing diverse human cell types, (iii) a pan-mammalian dataset (GSE223748) spanning dozens of species, and (iv) a brain single-cell methylation dataset generated using the recent sciMETv3 method (GSE273592) [30]. We also finetuned the larger model for the mammalian and single-cell datasets, given they are outside of the training domain of human bulk methylation. For qualitative and quantitative analyses respectively, we used UMAP to reduce these high-dimensional embeddings to two dimensions, and we quantified clustering quality using ARI and NMI (Supplementary Table 5). In the AltumAge dataset (Figure 2c), distinct clusters corresponding to tissues such as placenta, brain, and blood emerged; CpGPT_100M_ embeddings achieved substantially higher clustering metrics (ARI = 0.234, NMI = 0.509) compared to raw beta values (ARI = 0.031, NMI = 0.083), confirming that CpGPT captures tissue-specific methylation signatures (Supplementary Figure 4). Similarly, in the RRBS atlas, samples mostly grouped by cell type, with separations observed among divergent lineages, and CpGPT_100M_ embeddings yielded the highest ARI (0.091) and NMI (0.332), with ARI improving from 0.001 and NMI from 0.146 over raw beta values (Supplementary Figures 5, 6). In the pan-mammalian dataset (Figure 2d, Supplementary Figure 7), the sample embeddings clustered by species, with species phylogenetically closer to Homo sapiens positioned in closer proximity; CpGPT_100M_ achieved the strongest species-level clustering (ARI = 0.606, NMI = 0.784), indicating that species identity is the dominant organizing principle in the pretrained embeddings. After finetuning on the mammalian dataset, the CpGPT_100M_-Mammalian variant shifted the embedding structure so that tissue type became the primary separator (tissue ARI = 0.322, NMI = 0.546), while species-level clustering decreased (ARI = 0.397 vs 0.606 pretrained). In the single-cell dataset, the finetuned (CpGPT_100M_-sciMETv3) sample embeddings clearly separated major brain cell populations, as confirmed by coloring cells according to independently derived annotations from the original sciMETv3 atlas [30] (Supplementary Figure 8). Quantitatively, CpGPT_100M_-sciMETv3 achieved the highest clustering metrics over raw beta values (ARI = 0.080 vs −0.012, NMI = 0.179 vs 0.026). In contrast, UMAP projections of the raw methylation values failed to resolve these populations, producing a largely undifferentiated embedding space. These observations demonstrate that CpGPT’s sample embeddings accurately reflect cellular identity, tissue type, and evolutionary relationships even in the absence of explicit labels.

An important application of robust sample embeddings is the ability to perform zero-shot reference mapping, where labels from a well-annotated reference dataset can be transferred to an unlabeled dataset without additional training. To evaluate this capability, we projected the CpGPT_100M_ sample embeddings of the Hannum dataset (GSE40279) [10] onto those from the AltumAge dataset, which contains comprehensive tissue-specific annotations (Figure 2e). Our objective was to determine whether the blood-derived samples from the Hannum cohort could be accurately classified into the tissue categories defined by AltumAge. The mapping results were compelling: out of 656 Hannum samples, 340 were classified as “blood leukocyte,” 230 as “blood buffy coat,” 83 as “blood whole,” and 3 as “other” (Figures 2f, 2g). To provide a more rigorous evaluation in addition to the Hannum cohort, which consists exclusively of blood samples, we applied the same zero-shot label-transfer protocol to another independent dataset of paired blood and brain samples from 20 Parkinson’s disease patients (GSE151355). In this setup, all 20 blood samples were correctly mapped to “blood whole,” and 17 of 20 brain samples were mapped to “brain frontal cortex” (Supplementary Figure 9). The three remaining brain samples were assigned to the “other” category. These results confirm that CpGPT_100M_’s sample embeddings capture specific signatures sufficient to distinguish tissues in a zero-shot manner.

Finally, to further demonstrate the practical utility of CpGPT’s sample embeddings, we applied zero-shot reference mapping to a dataset capturing the reprogramming of fibroblasts to pluripotency (GSE54848) [31]. Using the comprehensive CpGCorpus as the reference, we mapped each sample from the reprogramming time course to its most similar entry in the Gene Expression Omnibus (Supplementary Table 6). The three replicates of day 0 fibroblasts (prior to OSKM induction) mapped to GSM868011, GSM868013, and GSM868015, corresponding to “somatic primary HDFs.” By day 28, the replicates mapped to GSM867971, GSM867983, and GSM867975, now annotated as “induced pluripotency stem cell.” Day 15 was the last time point where the mapped embeddings retained fibroblast descriptions, in agreement with previous research that identifies this period as the transition point when fibroblasts lose their cellular identity [32]. This example demonstrates that CpGPT’s sample embeddings are not only biologically informative but also robust enough to capture dynamic changes in cell state during reprogramming.

### 2.3 CpGPT faithfully reconstructs the methylome

DNA methylation data often contain missing values or unmeasured CpG sites because of platform differences and variations in experimental design. An ideal foundation model should be able to leverage its learned knowledge to fill in these gaps and infer the methylation status of CpG sites that were never directly observed. This ability, often referred to as zero-shot imputation, is particularly valuable for applications like biomarker discovery, epigenetic clock predictions, and data integration across different methylation arrays.

Here, we systematically evaluated CpGPT’s capacity to perform zero-shot imputation and array conversion in several scenarios, including reconstructing sites omitted during pretraining, converting between Illumina arrays, mapping methylation patterns across mammalian species, and even handling single-cell methylation data. We compare CpGPT against four statistical baselines: the sample mean (the per-sample global mean methylation across all observed probes), the sample chromosome mean (the per-sample mean methylation within each chromosome), the population chromosome mean (the training-set pooled mean within each chromosome), and the population locus mean (the per-probe training-set mean).

In the first experiment, our objective was to assess the generality of CpGPT’s imputation when it encounters CpG sites that were deliberately excluded from its pretraining (Supplementary Tables 7, 8). Specifically, we removed 1% of CpG sites during model training and then tested how well CpGPT could infer their methylation levels in the CpGCorpus test set (Figure 3a). In the simplest case (“seen, seen”), the model receives beta values from CpG sites it had encountered during pretraining and predicts the methylation of target loci also within the pretraining universe. Both CpGPT_2M_ and CpGPT_100M_ outperformed all baselines (sample mean MAE = 0.3312 ± 0.0374; sample chromosome mean MAE = 0.3295 ± 0.0375; population chromosome mean MAE = 0.3530 ± 0.0305; population locus mean MAE = 0.3947 ± 0.0362; CpGPT_2M_ MAE = 0.0726 ± 0.0244; CpGPT_100M_ MAE = 0.0386 ± 0.0218). We then tested the more challenging “seen, unseen” case, in which the model predicts completely new loci that were never observed in training. Again, both CpGPT models substantially outperformed every baseline (sample mean MAE = 0.3327 ± 0.0366; sample chromosome mean MAE = 0.3276 ± 0.0401; population chromosome mean MAE = 0.3537 ± 0.0297; CpGPT_2M_ MAE = 0.1529 ± 0.0261; CpGPT_100M_ MAE = 0.0723 ± 0.0222). Even in the most stringent “unseen, unseen” setup, where both the input and predicted sites were not in the pretraining universe, CpGPT beat all baselines (sample mean MAE = 0.3329 ± 0.0366; sample chromosome mean MAE = 0.3330 ± 0.0348; population chromosome mean MAE = 0.3537 ± 0.0300; CpGPT_2M_ MAE = 0.1457 ± 0.0226; CpGPT_100M_ MAE = 0.0811 ± 0.0247). To assess whether performance varies by tissue type, we stratified the CpGCorpus test set into 10 tissue categories using GEO metadata (Supplementary Table 9). CpGPT_100M_ consistently outperformed all baselines across tissues in the “unseen, unseen” setup, with the lowest MAE in blood and immune cell samples (0.068 and 0.069, respectively) and the highest in fibroblasts/skin (0.122). These outcomes show that CpGPT generalizes to novel genomic regions purely from previously learned representations.

**Figure 3:**
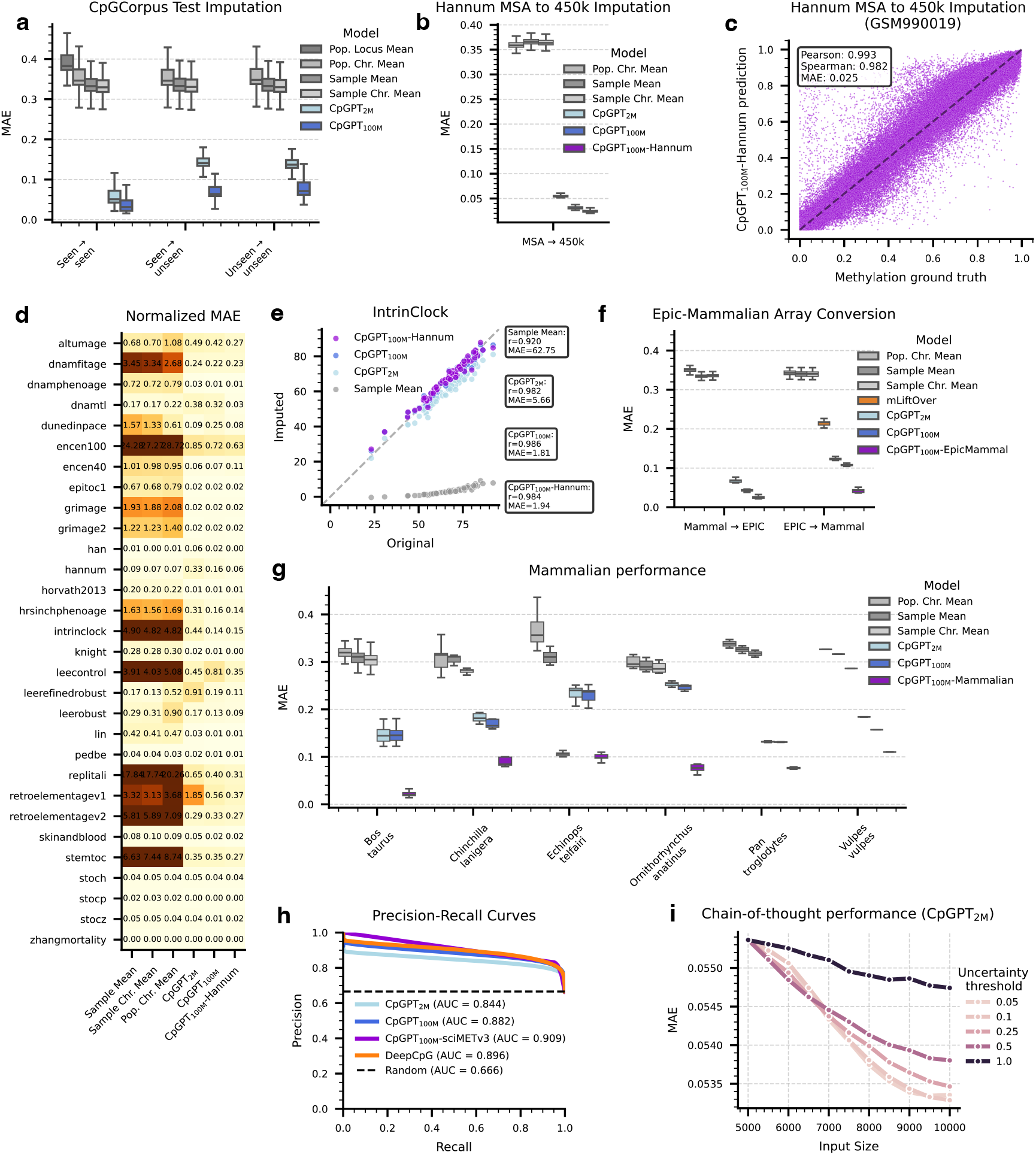
CpGPT performs zero-shot imputation across various conditions. (a) Boxplot of average mean absolute error (MAE) per sample for CpGPT_2M_, CpGPT_100M_, and statistical baselines, tested on the CpGCorpus dataset. The 5,000 input CpG sites and the 5,000 queried CpG sites are either from the pretraining vocabulary (“seen”) or not (“unseen”). Boxes show the first and third quartiles (IQR), whiskers extend to the last data point within 1.5 IQR, and the middle line marks the median. (b) Boxplot of the average MAE per sample in the Hannum dataset for statistical baselines, CpGPT_2M_, CpGPT_100M_, and finetuned CpGPT_100M_-Hannum models. The query includes all CpG sites in the 450k array not present in the MSA array. Boxes show the IQR, whiskers extend to the last data point within 1.5 IQR, and the middle line marks the median. (c) Scatterplot comparing the ground-truth beta values versus CpGPT_100M_-Hannum predictions for sample GSM990019, showing the conversion from MSA probes to remaining 450k probes. (d) Heatmap of the MAE between epigenetic clock calculations using only MSA-overlapping probes (zeroed-out missing probes) or CpGPT-imputed probes (CpGPT_2M_, CpGPT_100M_, finetuned CpGPT_100M_-Hannum), and statistical baselines, compared against the full 450k values (Hannum dataset). (e) Scatterplot comparing intrinclock predictions using only MSA-overlapping probes (missing probes either zeroed out or imputed with CpGPT_2M_, CpGPT_100M_, finetuned CpGPT_100M_-Hannum) or sample mean to the ground-truth 450k array (Hannum dataset). (f) Boxplot of the average MAE per sample for statistical baselines,mLiftOver, CpGPT_2M_, CpGPT_100M_, and finetuned CpGPT_100M_-EpicMammal when converting between paired EPIC and Mammalian40k arrays. mLiftOver only supports EPIC to Mammalian40k conversion. Results for Vulpes vulpes are from a single sample. Boxes show the IQR, whiskers extend to the last data point within 1.5 IQR, and the middle line marks the median. (G) Boxplot of the average MAE per sample for a statistical baselines, CpGPT_2M_, CpGPT_100M_, or finetuned CpGPT_100M_-Mammalian model. Results are shown for a random subset of 5,000 CpG sites in six unseen mammalian species. Error bars represent the standard deviation. (h) Precision-recall curves for statistical baselines (sample mean, chromosome means, locus mean), DeepCpG, CpGPT_2M_, CpGPT_100M_, and finetuned CpGPT_100M_-sciMETv3, showing imputation performance on single-cell methylation (sciMETv3 dataset). (i) Lineplot of the MAE for MSA-to-450k conversion (Hannum dataset) using CpGPT_2M_ with iterative refinement. Different uncertainty quantiles are used to filter generated CpG methylation values at each refinement step.

We next explored a more practical scenario of reconstructing methylation sites for an emerging array platform, the Illumina MSA [33], for which no paired MSA and conventional array data exist. We simulated this situation using the Hannum dataset (GSE40279) [10], originally measured on the Illumina 450k array. To mimic partial MSA data, we kept only the 113,585 probes shared between 450k and MSA, then used CpGPT to impute the remaining 450k probes needed for various epigenetic clocks (Figure 3b). Even without direct MSA-450k paired training data, CpGPT_100M_ produced an MAE of 0.0314 ± 0.0030, and a finetuned version (CpGPT_100M_-Hannum) achieved an MAE of 0.0244 ± 0.0029, both outperforming the sample mean (MAE = 0.3661 ± 0.0071). As shown by the near-perfect identity line fit for an example sample (Figure 3c), the model accurately recovered large segments of the 450k methylome from a subset of MSA-like inputs.

Because epigenetic clocks often rely on sets of probes that may be missing from a given platform, we then tested the downstream impact of CpGPT-based array conversion on several clocks, including *encen40* [34], *intrinclock* [35], and *stemtoc* [36]. We computed these clocks on the Hannum dataset using fully measured 450k data (considered the ground truth) and compared them to predictions derived from CpGPT-imputed values with a subset of the MSA probes as input (Figure 3d; Supplementary Figure 10). The clock estimates from CpGPT-imputed data closely matched those from the fully measured set. For instance, the correlation of the *intrinclock* with chronological age improved from 0.920 (sample mean imputation) to 0.984 when relying on CpGPT_100M_-Hannum predictions (Figure 3e). These results demonstrate that CpGPT’s array conversion can restore missing probes in a manner that preserves or even enhances the performance of modern epigenetic clocks.

We next aimed to convert data between the Illumina EPIC and Horvath mammalian arrays [37]. Using paired blood samples, we assessed two directions: reconstructing EPIC from the mammalian array and vice versa. As illustrated in Figure 3f, for conversion to EPIC, CpGPT_100M_-EpicMammal yielded an MAE of 0.0251 ± 0.0030 versus a sample mean of 0.3349 ± 0.0062, whereas for conversion to the mammalian array, it achieved 0.0419 ± 0.0043 versus a sample mean of 0.3406 ± 0.0081. We also compared against mLiftOver [38], a rule-based tool that allows the conversion of blood samples from EPIC to the mammalian array (mLiftOver does not support the reverse), mLiftOver achieved an MAE of 0.2164 ± 0.0107, compared to 0.0419 ± 0.0043 for CpGPT_100M_-EpicMammal. Despite the lower fidelity of the epigenetic clock predictions from the reconstructed data (Supplementary Figures 11, 12), these findings show that CpGPT is capable of bridging even larger design gaps between array platforms, thereby unlocking cross-platform analyses such as applying mammalian-focused epigenetic clocks to EPIC data. Moreover, unlike tissue-specific methods such as *mLiftOver* [38], CpGPT’s multi-tissue training in *CpGCorpus* enables tissue-agnostic imputation, making it more broadly applicable.

We further tested CpGPT’s ability to generalize across species by using data from 55 mammalian taxa (GSE223748) [39, 40]. From these, 43 species comprised the training set, 6 formed a validation set, and 6 served as an unseen test set. Within each species, half of the CpG sites were masked, and CpGPT was tasked to impute those sites (Figure 3g). For a species it had never seen during training, Pan troglodytes (chimpanzee), CpGPT_100M_-Mammalian achieved an MAE of 0.0814 ± 0.0036, compared to a sample mean of 0.3257 ± 0.0131. Although performance tended to be stronger for species phylogenetically closer to humans, these results reveal that a model solely trained on human arrays can still capture salient epigenetic patterns in evolutionarily related mammals. This cross-species flexibility opens the door to unified frameworks for translational studies and cross-mammalian biomarker research.

Next, we tested if CpGPT could handle single-cell methylation data, which are typically sparse and largely binary. We used the sciMETv3 dataset (GSE273592) [30], focusing on the area under precision–recall (AUPRC) to evaluate its predictions (Figure 3h). In addition to the statistical baselines, we compared CpGPT to DeepCpG [41], a method specifically designed for single-cell methylation imputation. All statistical baselines achieved near-random AUPRC, confirming the non-trivial nature of single-cell imputation. CpGPT_100M_ reached an AUPRC of 0.882, whereas the finetuned CpGPT_100M_-sciMETv3 scored 0.909. DeepCpG, whilst performing better than the zero-shot CpGPT model, lagged slightly behind the fine-tuned version with an AUPRC of 0.897. On the area under the receiver-operator curve (AUROC), DeepCpG led, with CpGPT_100M_ second followed by CpGPT_100M_-sciMETv3 and CpGPT_2M_-sciMETv3 (Supplementary Figure 13). Overall, CpGPT performs strongly for single-cell DNA methylation imputation.

Finally, we investigated whether CpGPT could benefit from iterative refinement at inference time, a strategy that leverages increased test-time computation to progressively improve predictions, akin to chain-of-thought in natural language [42]. In this procedure, we first provided the model with a set of known CpG sites and generated methylation predictions for a random subset of unmeasured positions. We then selected the most confident predictions (those with the lowest estimated uncertainty) and appended them to the input, allowing the model to refine its estimates in subsequent iterations. To evaluate this strategy, we used the Hannum dataset’s array-conversion task as a test bed (Figure 3i). Starting with 5,000 MSA CpG sites and CpGPT_2M_, the mean absolute error (MAE) decreased from 0.0554 ± 0.0023 to 0.0534 ± 0.0023 after 10 refinement steps of size 500 under the 5th-percentile uncertainty threshold (best MAE = 0.0533 at 9 steps; full results in Supplementary Table 10). We also examined how larger step sizes (1,000 or 2,000) and finetuning influenced the iterative refinement curve, finding that stricter uncertainty filtering improved accuracy while step size exerted minimal effect (Supplementary Figures 14, 15). However, beyond a certain number of iterations, performance gains deteriorated, likely because accumulated prediction noise begins to outweigh the benefit of additional context [43]. These results demonstrate that CpGPT’s framework can be adapted with a chain-of-thought approach to improve accuracy at test time.

### 2.4 CpGPT displays state-of-the-art performance when finetuned

Foundation models often gain substantial performance improvements when finetuned on specific tasks, because the pretraining process captures broad, generalizable representations that can be readily adapted to particular datasets. We hypothesized that this principle would extend to CpGPT, enabling the model to excel in a variety of downstream prediction challenges once it had been exposed to the unique nuances of each problem. To test this, we employed our lighter version, CpGPT_2M_, and examined its performance on (i) regression tasks of varying complexity, (ii) multiclass regression, and (iii) binary classification.

In the first experiment, we assessed whether CpGPT_2M_ could predict chronological age from multi-tissue data with greater accuracy than conventional methods. We partitioned the AltumAge dataset [16] into training, validation, and testing sets, and compared CpGPT_2M_-Age to an ElasticNet model and a multilayer perceptron (MLP) (Supplementary Table 11). In terms of mean squared error (MSE) and MAE, CpGPT_2M_-Age achieved an MSE of 38.13 years^2^ (MAE = 3.29 years), outperforming both ElasticNet (MSE = 270.4, MAE = 11.37) and MLP (MSE = 86.56, MAE = 5.51). Moreover, the Pearson correlation coefficient rose to 0.9847 with CpGPT_2M_-Age compared to 0.9116 and 0.9679 for ElasticNet and MLP, respectively (Figure 4a). These results suggest that incorporating sequence and epigenetic context through CpGPT yields a more nuanced representation of methylation data, ultimately enhancing the accuracy of age estimation.

**Figure 4:**
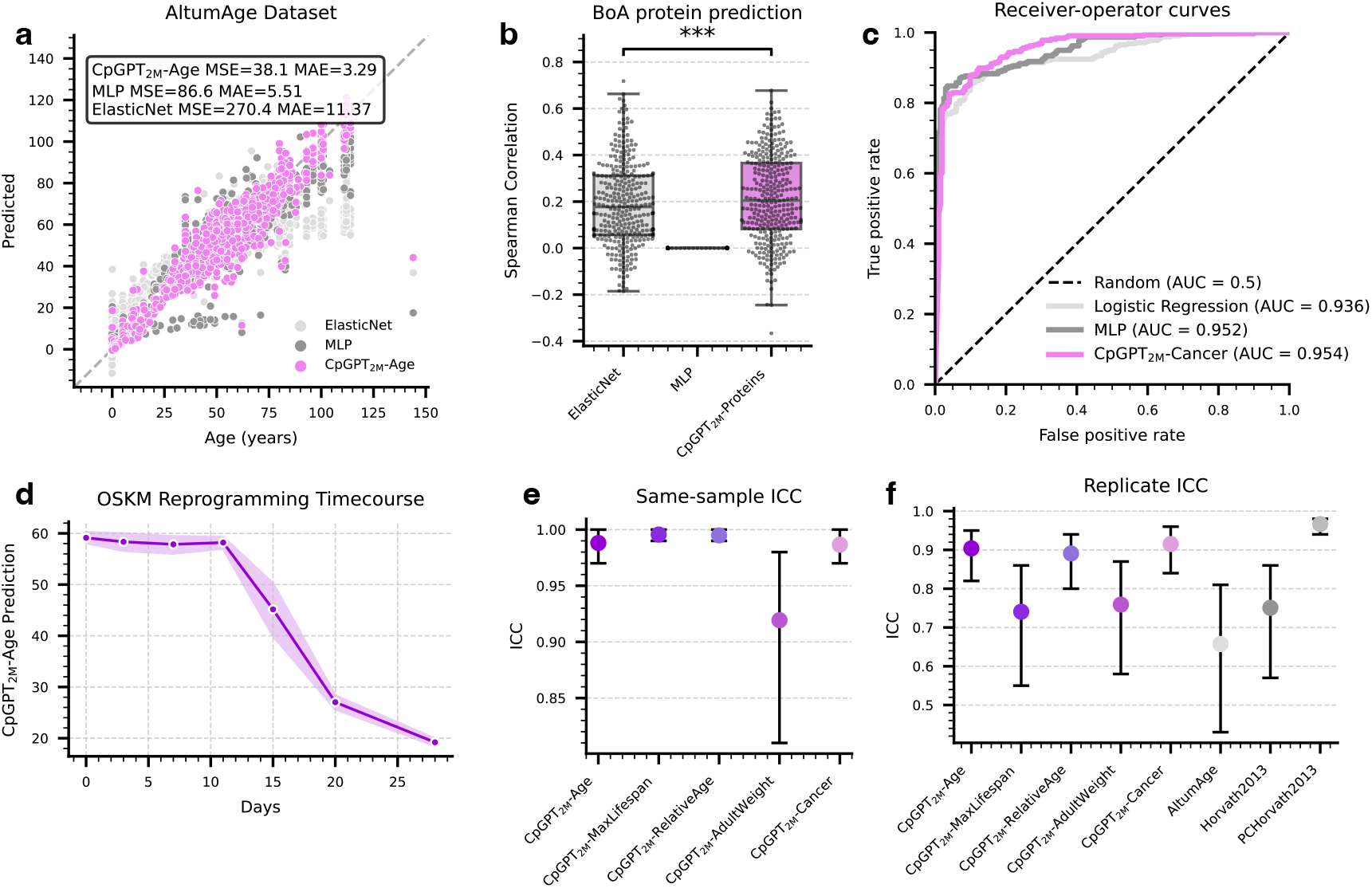
CpGPT outperforms neural networks and linear models, while showing reproducible results. (a) Scatterplot comparing ground-truth age to predictions on the AltumAge dataset using a linear model (ElasticNet), a multi-layer perceptron (MLP), and CpGPT_2M_-Age. (b) Boxplot of the Spearman correlation between ground-truth and predicted blood protein levels for 322 proteins in the Biomarkers of Aging dataset (GSE246337), comparing ElasticNet, MLP, and CpGPT_2M_-Proteins. (c) Receiver operating characteristic (ROC) curves of cancer prediction on the cancer dataset for a random baseline, logistic regression, MLP, and CpGPT_2M_-Cancer. (d) Lineplot (with 95% confidence intervals) of CpGPT_2M_-Age predictions over an OSKM reprogramming time course using random subsets of CpG sites. (e) Dotplot of the intraclass correlation coefficient (ICC) for all five finetuned CpGPT models with phenotype prediction heads (Age, MaxLifespan, RelativeAge, AdultWeight, Cancer) on the OSKM reprogramming dataset, computed over 10 random CpG subset inputs. Error bars show the 95% confidence intervals. (f) Dotplot of the ICC for various CpGPT models and three epigenetic clocks on a reliability dataset with technical replicates, all using a fixed input subset of 10,000 CpG sites.

We then investigated a more demanding multiclass regression problem involving proteomic measurements. Here, our goal was to predict the levels of 322 proteins that were quantified alongside methylation data for the Biomarkers of Aging Consortium (GSE246337) [44]. This task is especially challenging because proteomic assays can be prone to measurement variability. Likely due to this potential noise, CpGPT_2M_-Proteins attained a median Spearman correlation coefficient of 0.204 across all proteins (median MSE = 0.868, MAE = 0.694), surpassing both ElasticNet (Spearman = 0.177, MSE = 0.898, MAE = 0.712) and MLP (Spearman = 0.0 due to mode collapse) (Figure 4b, Supplementary Figure 16, Supplementary Table 12). These findings underscore CpGPT’s performance when predicting hundreds of variables with a single forward pass.

We next applied CpGPT to a binary classification task aimed at distinguishing between normal and adjacent cancerous tissues, using a dataset from multiple studies [16]. When comparing precision–recall and receiver operating characteristic curves (Figure 4c and Supplementary Figure 17), CpGPT_2M_-Cancer achieved an AUROC of 0.9544 and AUPRC of 0.9651, closely matching the MLP (AUROC 0.9519, AUPRC 0.9722) and outperforming logistic regression (AUROC 0.9357, AUPRC 0.9646). Thus, CpGPT not only excels at regression tasks but also maintains robust classification performance.

We further examined how CpGPT would handle data beyond human samples by finetuning on a mammalian dataset to predict relative age, adult bodyweight, and maximum lifespan. The model exhibited lower error compared to ElasticNet and MLP baselines across all three tasks: relative age (CpGPT_2M_-RelativeAge MAE = 0.143, MSE = 0.029 vs. ElasticNet MAE = 0.158, MSE = 0.030), average adult weight (CpGPT_2M_-AdultWeight MAE = 2.66, MSE = 7.32 vs. ElasticNet MAE = 3.99, MSE = 16.70), and maximum lifespan (CpGPT_2M_-MaximumLifespan MAE = 0.151, MSE = 0.055 vs. ElasticNet MAE = 0.643, MSE = 0.437), suggesting that the pretrained embeddings encode relationships that generalize across different mammalian species (Supplementary Figures 18, 19, 20). Although the performance was not as strong as in the age prediction or cancer classification tasks, these results highlight the flexibility of the framework.

To evaluate the consistency of CpGPT’s predictions when given different subsets of methylation data, we analyzed the OSKM dataset (GSE54848) with 10 distinct random seeds to select different subsets of 10,000 probes from the Illumina 450k array. We specifically tracked how CpGPT_2M_-Age behaves throughout a reprogramming time course, alongside other finetuned models (Supplementary Figure 21). As shown in Figure 4d, the predicted chronological age declined over time, reflecting the dedifferentiation process, and only the adult bodyweight predictor exhibited a slight upward drift. To quantify stability, we computed the intraclass correlation coefficient (ICC) across predictions from different subsets of CpG sites for each finetuned model (Figure 4e). Most predictors had ICC values above 0.99, and the lowest (CpGPT_2M_-AdultWeight) reached 0.919. These high ICCs illustrate that CpGPT remains consistent even when the input methylation sites are changed, affirming its reliability in scenarios where coverage or probe availability may vary.

Finally, we benchmarked CpGPT’s stability against replicate samples in a blood methylation dataset (GSE55763) [45]. We compared CpGPT predictions with three well-known multi-tissue epigenetic clocks (AltumAge [16], Horvath2013 [9], and PCHorvath2013 [14]), fixing the input to a randomly selected subset of 10,000 CpG sites (Figure 4f). Although PCHorvath2013 exhibited a higher ICC (0.967) than CpGPT_2M_-Age (0.904), the dynamic range of PCHorvath2013’s predictions was small (only about 0.7 years), whereas CpGPT_2M_-Age spanned approximately 12 years (Supplementary Figure 22). As a result, CpGPT achieved a lower mean squared error and mean absolute error (MSE = 86.42, MAE = 7.38) relative to these clocks (AltumAge MSE = 362.89, MAE = 16.59; Horvath2013 MSE = 393.20, MAE = 17.54; PCHorvath2013 MSE = 136.50, MAE = 9.58), indicating that the model not only preserves reproducibility but also captures a wider range of variation in biological age estimates from a small subset of the methylome. This combination of accuracy, robustness, and generalizability underscores the advantages of finetuning CpGPT for specific tasks, paving the way for more precise and flexible analyses of methylation data

### 2.5 CpGPT strongly predicts mortality and morbidity

To showcase the downstream utility of finetuned CpGPT predictions, we developed a mortality predictor named CpGPTGrimAge3 based on the CpGPT_2M_-Proteins model. Specifically, we adopted GrimAge’s methodology by first predicting protein proxies from methylation data and then applying a Cox proportional hazards model to predict mortality [11, 12] (see Methods). CpGPTGrimAge3 was trained on the Framingham Heart Study (FHS) cohort (N = 3,853; deaths n = 319) and tested on three independent holdout cohorts: MGB (N = 4,387; deaths n = 739), WHI (N = 2,101; deaths n = 563), and BLSA (N = 768; deaths n = 296). We benchmarked CpGPTGrimAge3 against eight established epigenetic clocks—PCHorvath1, PCHannum, PhenoAge, PCPhenoAge, DunedinPACE, SystemsAge, GrimAgeV2, and PCGrimAge—across all cohorts.

For all-cause mortality, we fit Cox proportional hazards models adjusting for age and reported hazard ratios (HRs) per standard deviation increase in clock score (Figure 5a, Supplementary Table 14). In the training cohort (FHS), CpGPTGrimAge3 achieved the second largest HR among all clocks. Critically, this superiority generalized to all three held-out test cohorts, with CpGPTGrimAge3 point estimates consistently exceeding comparator clocks: approximately 3.7 (MGB), 2.1 (WHI), and 4.6 (BLSA). First-generation clocks such as PCHorvath1 and PCHannum showed markedly attenuated associations across all held out cohorts, while PCGrimAge and GrimAgeV2 performed comparably but remained below CpGPTGrimAge3 in most settings.

**Figure 5:**
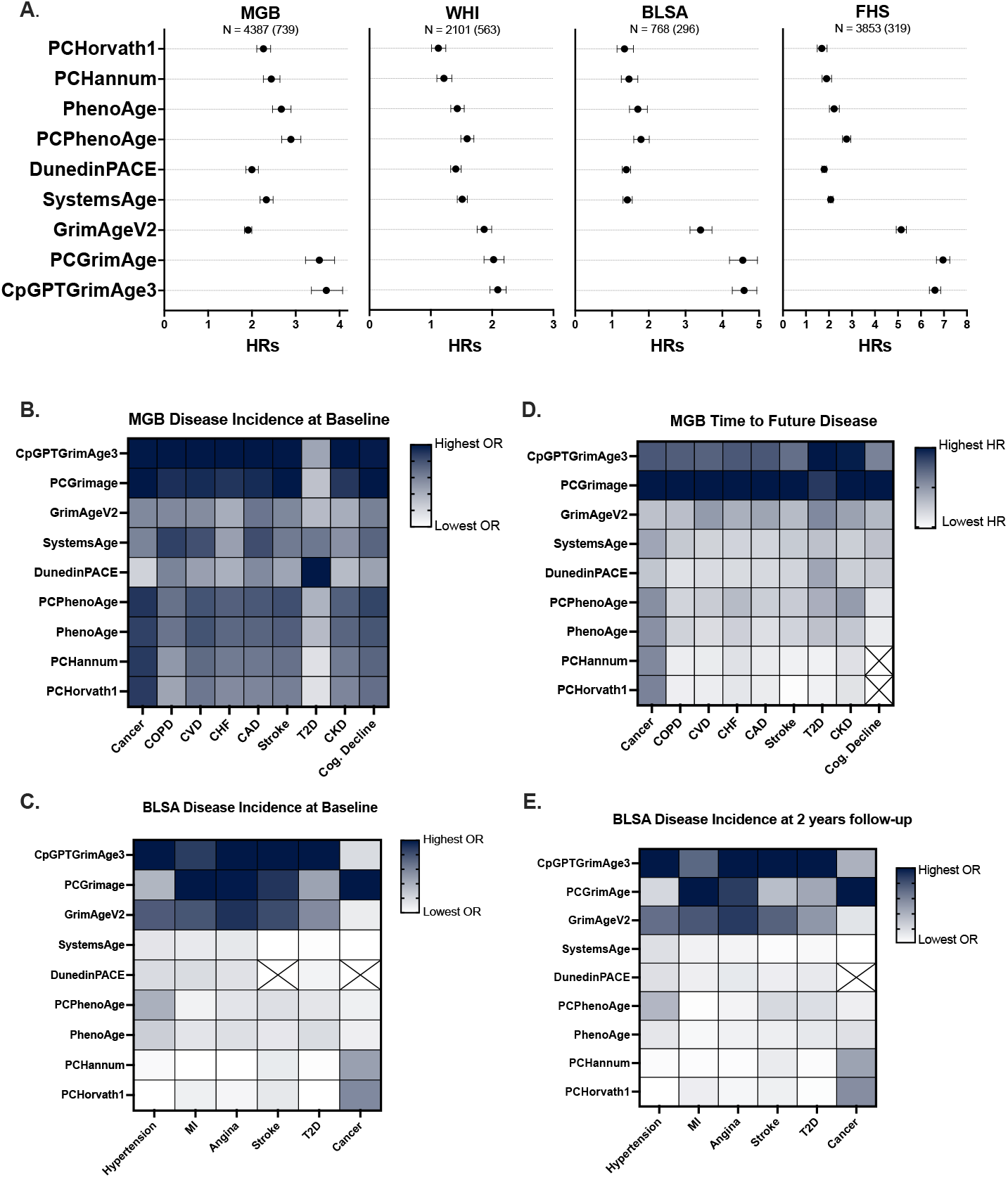
Predictive performance of CpGPTGrimAge3 and comparator epigenetic clocks for mortality and disease across independent cohorts. (A) Hazard ratios (HRs) per standard deviation increase in clock score for all-cause mortality across four cohorts: MGB (N = 4,387; 739 deaths), WHI (N = 2,101; 563 deaths), BLSA (N = 768; 296 deaths), and FHS (N = 3,853; 319 deaths), comparing CpGPTGrimAge3 against eight established clocks including PCHorvath1, PCHannum, PhenoAge, PCPhenoAge, DunedinPACE, SystemsAge, GrimAgeV2, and PCGrimAge. Points represent HR estimates and horizontal lines represent 95% confidence intervals. (B, C) Heatmaps depicting cross-sectional associations (odds ratios) between clock scores and prevalent disease at baseline in MGB (B; across cancer, COPD, CVD, CHF, CAD, stroke, T2D, CKD, and cognitive decline) and BLSA (C; across hypertension, myocardial infarction, angina, stroke, T2D, and cancer). (D) Heatmap of longitudinal associations (HRs) between clock scores and time to incident disease in MGB. (E) Heatmap of associations between clock scores and disease incidence at 2-year follow-up in BLSA. Across cohorts and outcomes, CpGPTGrimAge3 consistently ranks among the strongest clocks for mortality and disease, with PCGrimAge as its closest competitor. Darker shading indicates higher effect sizes; crossed cells indicate insufficient events for analysis.

Beyond mortality, we assessed associations between clock scores and prevalent disease at baseline across nine disease categories—cancer, COPD, CVD, CHF, CAD, Stroke, Type 2 Diabetes, CKD, and cognitive decline—in MGB (Figure 5b, Supplementary Table 15) and six disease categories—hypertension, MI, angina, stroke, T2D, and cancer—in BLSA (Figure 5c), using logistic regression models and reporting odds ratios. We further examined longitudinal associations with time to incident disease in MGB using Cox models (Figure 5d), and with disease incidence at two-year follow-up in BLSA (Figure 5e). Crossed cells indicate disease categories with insufficient events for stable estimation.

Across both cross-sectional and longitudinal analyses in held-out cohorts, CpGPTGrimAge3 ranked among the top two clocks for the large majority of disease endpoints, with PCGrimAge as its closest competitor. In MGB baseline disease prevalence analyses, CpGPTGrimAge3 ranked first in 1 of 9 disease categories and second in the remaining 8 categories, achieving a top-two finish in all 9 categories. In BLSA baseline analyses, CpGPTGrimAge3 ranked first in 4 of 6 disease categories and achieved a top-two finish in 5 of 6 categories. For incident disease prediction in MGB, CpGPTGrimAge3 ranked first in 6 of 9 disease categories and second in 2 additional categories, corresponding to a top-two finish in 8 of 9 categories. In BLSA two-year follow-up analyses, CpGPTGrimAge3 ranked first in 4 of 6 disease categories and achieved a top-two finish in 4 of 6 categories. Overall, across all five evaluation panels, CpGPTGrimAge3 achieved a top-two ranking in 30 of 34 clock–outcome comparisons (88.2%) and ranked first in 18 of 34 comparisons (52.9%). The consistency of these results across MGB, WHI, and BLSA, none of which contributed to training, provides strong evidence that CpGPTGrimAge3 generalizes robustly across diverse, independent populations, underscoring its utility in both research and clinical contexts.

## 3 Discussion

In this study, we introduce CpGPT, a foundation model specifically designed for DNA methylation analysis. By leveraging *CpGCorpus*, a harmonized resource comprising over 150,000 samples from diverse tissues and conditions, CpGPT captures relationships in the methylome across multiple levels of organization. Our results show that the model learns informative representations at both the CpG site and sample levels, and that these representations are useful across a broad range of tasks, including zero-shot imputation, array conversion, reference mapping, and supervised prediction of age, mortality, and morbidity.

The development of CpGPT addresses a significant gap in DNA methylation research. Traditional epigenetic clocks and methylation analyses often rely on linear models over fixed sets of CpG sites, approaches that have been highly useful but may not fully capture the higher-order dependencies present in the methylome [46, 16]. Although newer two-layer explainable perceptron models for DNA methylation to predict mortality have been developed, they still do not model the broader contextual structure of the aging methylome [47, 48]. By adopting a transformer architecture [1], CpGPT integrates sequence context, genomic position, and methylation state within a single framework. What is particularly encouraging is that these learned representations aligned with functional genomic annotations such as chromatin states and CpG islands without explicit supervision. Likewise, the sample embeddings organized according to tissue identity, cell type, and phylogenetic relatedness. Taken together, these findings suggest that CpGPT is learning biologically meaningful structure rather than simply memorizing probe-level correlations.

One of the most notable strengths of CpGPT is its ability to generalize across datasets and experimental settings. The zero-shot reference mapping of the Hannum [10] cohort onto AltumAge [16] tissue annotations, as well as the accurate transfer of labels in an independent paired blood-brain dataset, shows that the sample embeddings retain information sufficient for tissue-level classification without additional training. This same generalization was evident in the imputation results. CpGPT reconstructed loci withheld during pretraining, recovered 450k probes from MSA-like inputs [33], converted between EPIC and mammalian arrays, generalized to unseen mammalian species, and performed strongly on sparse single-cell methylation data. These are practical settings that routinely limit downstream methylation analyses, and they highlight an important advantage of a foundation model approach: the same pretrained representation can be repurposed for harmonization, annotation, and cross-platform integration. The modest gains observed with iterative refinement at inference time further suggest that test-time compute may be another useful axis for improving methylation prediction.

When finetuned, CpGPT also performed strongly across a range of downstream tasks, including chronological age prediction, proteomic proxy prediction, cancer classification, and mammalian trait prediction. In the aging context, the mortality model CpGPTGrimAge3 was especially notable. It achieved the strongest age-adjusted association with mortality in all three independent test cohorts (MGB, WHI, and BLSA) and the second-strongest in the FHS training cohort, behind only PCGrimAge, consistently outperforming established first- and second-generation epigenetic clocks. Beyond mortality prediction, CpGPTGrimAge3 demonstrated strong and highly consistent associations with both prevalent and incident disease outcomes across multiple independent cohorts. Across 34 mortality and disease endpoints evaluated, CpGPTGrimAge3 ranked among the top two performing clocks in 30 comparisons (88.2%) and ranked first in 18 (52.9%), highlighting its broad predictive utility across diverse aging-related phenotypes.

At the same time, several limitations remain. Our pretraining data are dominated by human bulk array-based methylation profiles, and although the model generalized surprisingly well beyond that setting, performance still varied across tissues, platforms, and species. In addition, specialized methods remained competitive in some domains, particularly for single-cell methylation, underscoring that foundation models complement rather than eliminate task-specific approaches. Future work should expand *CpGCorpus* beyond array data, incorporate broader non-human and longitudinal datasets, and further explore how uncertainty estimates and iterative inference can improve reliability and interpretability.

Overall, we view CpGPT as a step toward a more general representation-learning framework for the methylome. Rather than building a separate model for each assay, tissue, or aging outcome, it may be possible to start from a shared epigenetic foundation model and adapt it to the biological question at hand. For aging research in particular, this is appealing because the biology of aging is distributed, context dependent, and only partly captured by fixed linear signatures. CpGPT offers a practical way to work with that complexity while opening new opportunities for biological discovery across health, disease, and longevity.

## 4 Methods

### 4.1 Data curation

#### 4.1.1 CpGCorpus

To train an effective foundation model capable of generalizing across diverse biological contexts, we curated a comprehensive DNA methylation database named *CpGCorpus*. This database aggregates publicly available DNA methylation data from the Gene Expression Omnibus (GEO) [49], encompassing a total of 2,042 studies, 155,151 human samples, and over 101 billion tokens (where each token is one CpG measurement in one sample). These samples were measured using various Illumina methylation array platforms, including the 27k, 450k, EPIC, EPIC+, and EPICv2 arrays, providing a wide spectrum of coverage across the human methylome. Across all platforms combined, CpGCorpus covers 1,174,502 unique CpG probes. The number of CpGs per sample varies by platform, ranging from ~27,000 for 27k arrays to ~860,000 for EPIC arrays. The collected data represent a rich diversity of tissue types, developmental stages, disease conditions, and demographic backgrounds. This diversity is crucial for training a foundation model that can capture the complex patterns and variations in DNA methylation across different biological states.

To ensure consistency and quality across the dataset, we performed the following:

- **Processing of raw data:** For datasets where raw IDAT files were available, we utilized the R package *SeSAMe* [50] for data processing. *SeSAMe* offers advanced normalization and preprocessing algorithms tailored for Illumina methylation arrays, including background correction, dye bias correction, detection p-value computation, and beta value estimation.
- **Handling of processed data:** For datasets lacking raw IDAT files, we employed the normalized beta value matrices provided by the original studies. These beta values were assumed to be preprocessed according to the methodologies described in their respective publications.
- **Quality control:** We implemented quality control measures to identify and exclude poor-quality samples and probes (*SeSAMe* argument `prep=“QCDPB”`). This included filtering out probes with detection p-values above a threshold, probes targeting single-nucleotide polymorphisms (SNPs), and those known to cross-hybridize. For data that was already processed, we excluded datasets from which beta values were outside of the expected 0 to 1 range.
- **Probe harmonization:** To address differences in probe sets across array platforms, we mapped probes to a common set of CpG sites based on their genomic coordinates (*SeSAMe* argument `collapseToPfx=TRUE`). This allowed us to integrate data from different arrays and ensured that the model learned from a consistent set of features.

The final *CpGCorpus* dataset provides a harmonized and high-quality resource for model training.

##### Data splits

To assess the performance and generalization capabilities of CpGPT, we partitioned *CpGCorpus* into training, validation, and test sets:

- **Training set:** Consists of 150,719 samples from 1,986 studies (median 30 samples per dataset). This set was used for model training. One percent of the available genomic locations were masked during pretraining.
- **Validation set:** Includes 174 samples from 10 studies (median 13 samples per dataset). This set was used for hyperparameter tuning and model selection.
- **Test set:** Contains 4,258 samples from 46 studies (median of 48 samples per study). This set was held during training and validation and used exclusively for the final evaluation of the model.

We ensured that there was no overlap of samples or studies between the splits to prevent data leakage. A detailed list of GEO Series (GSE) entries included in each split is provided in Supplementary Table 13.

#### 4.1.2 AltumAge

The AltumAge dataset was obtained from a public Google Drive repository [16]. This curated dataset includes age-labeled DNA methylation data from multiple studies, some of which make part of the training, validation or testing of *CpGCorpus*. We used a study-level splitting strategy, assigning 70% of the unique studies to training, 10% to validation, and 20% to testing.

#### 4.1.3 Cancer

The cancer dataset was downloaded from a public Google Drive repository [16]. It contains DNA methylation data from normal and adjacent cancerous tissue for multiple tissue types from several studies. Any beta values outside the 0–1 range were set to `NaN`. A study-level 50%-10%-40% train-val-test split was applied to create training, validation, and test sets.

#### 4.1.4 EPIC-Mammalian comparison

The EPIC-Mammalian dataset contains paired measurements from the EPIC and Horvath mammalian arrays for human blood samples. A 50%-10%-40% train-val-test split was used.

#### 4.1.5 Biomarkers of Aging Consortium

We accessed the Biomarkers of Aging Consortium (GSE246337) dataset via the *biolearn* Python package. Paired DNA methylation and protein measurements were taken from the same individuals, and proteins with duplicate measurements were averaged. For CpG site selection, we chose the superset of CpG sites used by GrimAge, GrimAge2, DunedinPACE, DNAmPhenoAge, and HRSInchPhenoAge [11, 12, 14, 13, 51]. The resulting dataset was divided into training (70%), validation (10%), and test (20%) splits.

#### 4.1.6 Mammalian

The Mammalian dataset, from GSE223748, captures DNA methylation data for over 100 mammalian species. However, we limited ourselves to the 55 species with the appropriate genomic mapping of the mammalian probes [52]. For data splits, human (*homo_sapiens*) and mouse (*mus_musculus*) data were retained in the training set; remaining species were split into training (80%), validation (10%), and test (10%). Two percent of the probes for training were reserved for out-of-distribution testing.

#### 4.1.7 Mortality

We utilized the Framingham Heart Study (FHS) cohort to develop CpGPTGrimAge3 and three independent cohorts—Massachusetts General Brigham (MGB), the Women’s Health Initiative (WHI), and the Baltimore Longitudinal Study of Aging (BLSA)—for held-out validation, with harmonized methylation data and clinical information in each. We excluded beta values outside the 0–1 range. The DNA methylation data from these studies were previously described [47, 48].

#### 4.1.8 OSKM

The OSKM dataset (GSE54848) includes 450k DNA methylation data from a reprogramming timecourse experiment of fibroblasts with cell sorting over a period of 28 days.

#### 4.1.9 Hannum

The Hannum dataset (GSE40279) contains blood DNA methylation data from the 450k array. The dataset was split with 80%-10%-10% for training, validation, and testing.

#### 4.1.10 RRBS atlas

The Human RRBS Atlas (GSE233417) captures methylation profiles using Reduced Representation Bisulfite Sequencing (RRBS). Original hg19 coordinates were converted to hg38, retaining only CpGs that correctly mapped to the updated reference and that overlapped the genomic loci from pretraining. We used a 50%-10%-40% train-val-test split. This ensured a balanced representation of tissue types across all subsets and facilitated tissue-specific analyses.

#### 4.1.11 sciMETv3

The sciMETv3 dataset (GSE273592) comprises single-cell methylation profiles generated with both Illumina and Ultima sequencers. We used the Illumina dataset. Only cells with at least 1,000 measured beta values from the genomic loci from pretraining were kept. The dataset was subsequently split into training (50%), validation (10%), and test (40%) sets.

#### 4.1.12 Reliability

The reliability dataset (GSE55763) was used to calculate intraclass correlation coefficients. The dataset comprises 36 whole blood samples with two technical replicates.

### 4.2 Model architecture

The CpGPT model is designed to integrate sequence, positional, and epigenetic information to effectively learn the complex relationships inherent in DNA methylation data. Below, we detail each component of the model architecture and the associated preprocessing steps.

#### 4.2.1 Input representation and preprocessing

The model input comprises DNA sequence embeddings, methylation beta values, and positional information for each CpG site. Specifically:

- **DNA sequence embeddings** (**E**): For each CpG site, we extracted a nucleotide sequence of length *L* centered on the target cytosine using the annotation from Ensembl version 112 [53]. These sequences were embedded into numerical representations using a pretrained DNA language model.
- **Methylation beta values** (*β*): The methylation level at each CpG site, represented as a beta value ranging from 0 (unmethylated) to 1 (fully methylated).
- **Positional information** (**p**): The genomic coordinates and chromosome indices for each CpG site.

To optimize the utilization of genomic context and mitigate potential biases, we applied the following preprocessing steps:

1. **Intra-chromosomal sorting:** Within each chromosome, CpG sites were sorted in ascending order based on their genomic coordinates. This preserves local genomic context and facilitates the modeling of spatial dependencies between neighboring CpG sites.
2. **Chromosome grouping:** CpG sites were grouped by chromosome to maintain proximate loci together.
3. **Stochastic chromosome shuffling:** The order of chromosomes was randomly shuffled for each input batch during training. Because each input is a variable-size subset of all CpG sites, a fixed chromosome order would allow the model to associate each input position with a specific CpG locus, learning positional shortcuts rather than genuine epigenetic dependencies. Shuffling ensures the model must rely on the sequence embeddings and positional encodings (which encode true genomic coordinates) rather than memorizing input order. Importantly, intra-chromosomal sorting is preserved, so local spatial relationships between neighboring CpG sites (including distance to telomeric or centromeric regions) are retained within each chromosome group.

This approach allows the model to capture both local (within chromosome) and global (across chromosomes) genomic contexts while remaining flexible to different input subsets and orderings at inference time.

#### 4.2.2 Sequence encoding

To encode the local genomic context of each CpG site, we employed a pretrained DNA language model to generate sequence embeddings:

- **Sequence extraction:** For each CpG site, a nucleotide sequence of length *L* centered on the CpG site was extracted.
- **Embedding generation:** These sequences were input into a DNA LLM [23, 20, 54], yielding embeddings 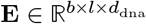, where *b* is the batch size, *l* is the sequence length, and *d*_dna_ is the dimension of the DNA embeddings.
- **Embedding transformation:** A trainable multi-layer perceptron (MLP) *f*_seq_ projected these embeddings into the model’s internal embedding space:

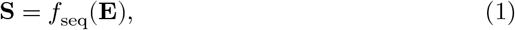

where 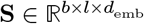 and *d*_emb_ is the model embedding dimension.

This process captures the local sequence context, enabling the model to learn patterns associated with sequence motifs and methylation patterns.

#### 4.2.3 Dual positional encoding strategy

We employed a dual positional encoding strategy that allows the model to uniquely identify each CpG location and understand CpG positions relative to each other:

##### Absolute positional encoding

We adapted the sinusoidal positional encoding scheme from Vaswani et al. [1] to suit genomic distances:

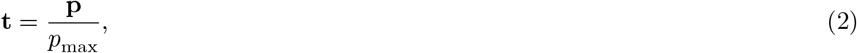

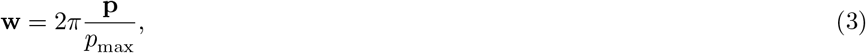

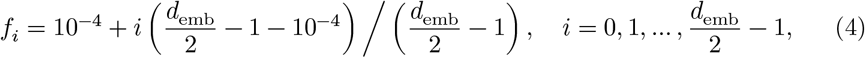

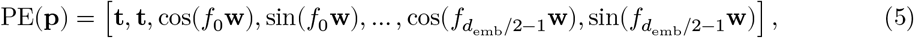

where *p*_max_ approximates the length in base pairs of the largest human chromosome (i.e., 3×10^8^ base pairs). This encoding PE(**p**) 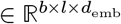 captures global positional information, which is needed for distinguishing between CpG sites with similar sequences but located in different genomic regions.

##### Relative positional encoding

To capture local positional relationships, we applied Rotary Positional Embeddings (RoPE) [21]:

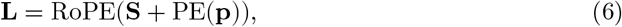

where 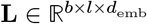 is the locus embedding. RoPE allows the model to incorporate relative positional information efficiently, enhancing its ability to model interactions between nearby CpG sites.

#### 4.2.4 Methylation state encoding

The methylation beta values were embedded to capture the epigenetic state:

- **Beta Value Embedding:** A trainable MLP *f*_*β*_ transformed the beta values into embeddings:

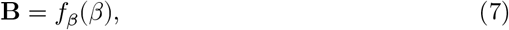

where 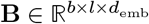.
- **CpG Site Embedding:** The final embedding for each CpG site combined the locus embedding and the beta value embedding:

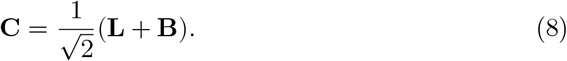

This normalization ensures that the combined embedding maintains a consistent scale.

#### 4.2.5 Transformer++ architecture

The core of CpGPT is based on the Transformer++ architecture [22], which enhances the original Transformer model [1] with:

1. **SwiGLU activation function:** Swish-Gated Linear Unit (SwiGLU) replaces the standard ReLU activation, providing smoother gradients and improved performance.
2. **RMSNorm pre-normalization:** Root Mean Square Layer Normalization stabilizes training in deep networks.
3. **Bias-free linear layers:** Removing bias terms reduces overfitting and parameter count.

The model consists of *N* layers of Transformer++ blocks. Each layer computes:

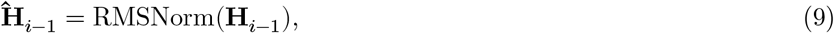

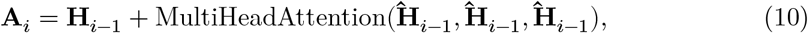

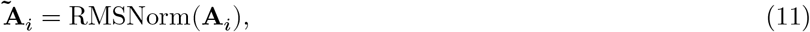

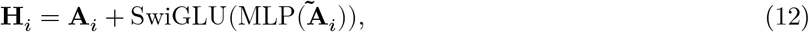

where **H**_0_ = [**c**_cls_; **C**], and 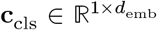 is a learnable classification token that aggregates information across the sequence.

#### 4.2.6 Decoders

We designed specialized decoders to extract meaningful outputs from the model:

##### M value decoder

Predicts methylation M values for CpG sites:

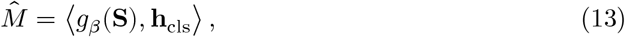

where *g*_*M*_ is a projection function, **h**_cls_ is the final hidden state of the classification token, and ⟨·, ·⟩ denotes the inner product. We opted for M values for the prediction as they have better statistical properties compared to beta values and there is no need for a sigmoid after the last hidden layer, which can cause gradient issues.

##### Uncertainty decoder

Estimates the uncertainty of the M value predictions:

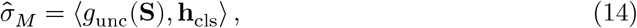

where *g*_unc_ is a projection function.

##### Condition decoder

Used during finetuning for specific downstream tasks:

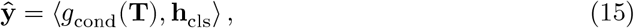

where **T** is a set of learnable condition tokens, and *g*_cond_ projects them into the embedding space.

### 4.3 Model variants

We pretrained two model variants and assigned each to the task regime where its capacity is best matched (Table 1). CpGPT_100M_ is used for all unsupervised tasks (embedding analysis and imputation), where the model must compress or reconstruct the full methylome. This high-dimensional reconstruction problem benefits directly from increased model capacity. CpGPT_2M_ is used for all supervised tasks (age regression, cancer classification, protein prediction, mortality), where the model maps methylation patterns to a single scalar or a small set of outputs. In this regime, task-specific supervision compensates for reduced model capacity.

**Table 1:** Comparison of CpGPT Model Versions.

| Parameter | CpGPT<sub>100M</sub> | CpGPT<sub>2M</sub> |
| --- | --- | --- |
| Total Parameters | 101M | 2.5M |
| d_embedding | 512 | 128 |
| d_hidden | 512 | 128 |
| n_attention_heads | 16 | 8 |
| n_layers | 32 | 8 |
| dropout | 0.01 | 0.01 |
| n_mlp_blocks | 3 | 2 |
| dna_context_len | 2001 | 2001 |
| max_length | 10000 | 5000 |

### 4.4 Pretraining procedure

#### 4.4.1 Training loop

We trained CpGPT using a multi-task learning approach. For each training batch, we performed the following steps:

1. **Data preparation:** A batch of samples *ℬ* = *β*, **E, c, p, O** was prepared, where **O** contains optional observed variables for downstream tasks.
2. **Missing value masking:** We created a mask **M**_na_ to handle missing beta values due to array differences or quality control filtering.
3. **Input-target split:** The beta values were randomly split into equally sized input and target sets using a 50% masking ratio:

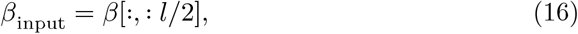

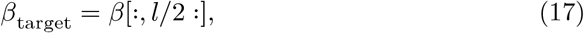

encouraging the model to reconstruct missing data. The choice of a 50% masking ratio balances two considerations. Compared with the 15% masking used in BERT [3], a higher ratio forces the model to learn more generalizable representations from limited input, which is critical for a reconstruction-oriented objective. Compared with the 75% masking used in Masked Autoencoders for vision [55], a lower ratio is appropriate here because individual CpG beta values carry more information per token than image patches (methylation values are continuous and less spatially redundant). Empirically, 50% masking also mirrors the practical deployment scenario in which CpGPT is used to impute substantial fractions of missing data across array platforms (e.g., converting Illumina 450k to EPIC, where roughly half the probes are absent).
4. **Encoding steps:**
  - Sequence encoding to obtain **S**.
  - Positional encoding to obtain **L**.
  - Methylation state encoding to obtain **B** and **C**.
5. **Transformer++ processing:** The combined embeddings were processed through the Transformer++ layers to obtain **H**_*N*_.
6. **Prediction:**
  - M value prediction 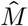.
  - Uncertainty estimation 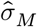.
  - Condition prediction ŷ (if applicable).

#### 4.4.2 Loss functions

The total loss *ℒ* is a weighted sum of several component losses:

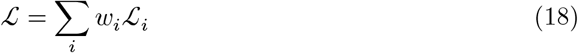

where *w*_*i*_ are the weights for each loss component. The main loss components are:

##### Methylation losses

A mean absolute error loss is used both for the predicted M values and the estimated error of the prediction itself. In addition, a Wasserstein distance loss, also known as Earth mover’s distance, is used to improve the distribution of the predicted M values when compared to the real distribution.

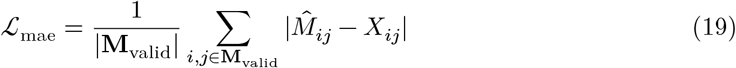

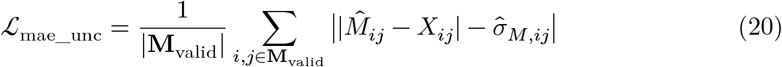

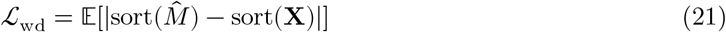

where **M**_valid_ = {non-NA indices in **X**_target_}

##### Sample embedding losses

To facilitate meaningful representations of the samples, the Kullback-Leibler divergence is incorporated to constrain the embedding space to be normally distributed with mean zero and variance one. Moreover, a contrastive loss first used by scGPT [4] is employed to separate dissimilar samples and bring together similar ones.

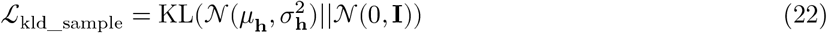

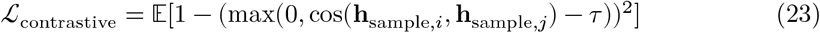

where *τ* is a threshold parameter, and 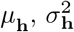 are the mean and variance of **h**_sample_ across the batch dimension.

##### Condition prediction loss (if enabled for finetuning)

The condition prediction loss *ℒ*_cond_ is task-specific and depends on the nature of the predicted conditions. Let ŷ ∈ ℝ^*b*×*k*^ be the predicted conditions and **O** ∈ ℝ^*b*×*k*^ be the observed variables, where *b* is the batch size and *k* is the number of conditions or features. The general form of the condition prediction loss is:

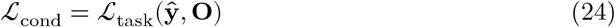

The specific form of *ℒ*_task_ depends on the prediction task:

1. For regression tasks:

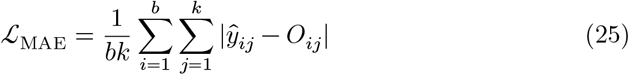

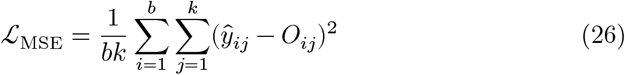
2. For binary classification tasks:

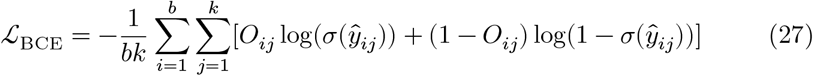

where *σ* is the sigmoid function.

#### 4.4.3 Software and hardware

CpGPT was developed and pretrained using *python* version 3.10.15 and *torch* version 2.4.1. A list of other dependencies will be available on our GitHub after publication.

The standard CpGPT model was trained on an NVIDIA H100 GPU for approximately 10 days.

### 4.5 Chain-of-thought inference

To enhance prediction accuracy during inference, we tested an iterative refinement procedure for CpGPT akin to chain-of-thought in natural language processing. This approach allows the model to progressively improve its methylation predictions by incorporating its own most confident estimates into subsequent iterations.

For a set of *n* refinement steps with a fixed step size *k*, the procedure operates as follows:

1. **Initialization:** Begin with an initial batch *ℬ*_0_ = {*β*_0_, **E**_0_, **c**_0_, **p**_0_} containing a small set of known CpG sites.
2. **Prediction generation:** For each refinement step *i* ∈ {1, 2, …, *n*}:
  - Calculate the candidate pool size 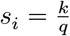, where *q* ∈ (0, 1) is the predetermined uncertainty quantile threshold.
  - Sample a random subset of *s*_*i*_ genomic locations 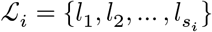 with available DNA LLM embeddings.
  - Extract DNA embeddings 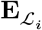 for these locations.
  - Generate methylation predictions and uncertainties:

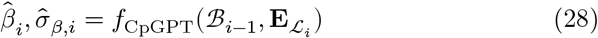
3. **Confidence-based filtering:** Compute the uncertainty threshold that selects approximately *k* predictions:

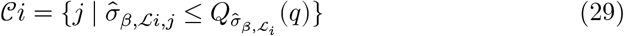

where 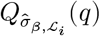 represents the *q*-th quantile of the uncertainty distribution 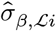. The size of *C*_*i*_ will be approximately *k* sites, ensuring a consistent step size across iterations.
4. **Batch augmentation:** Update the batch with confident predictions for the next iteration:

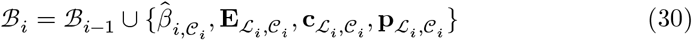
5. **Sorting:** Reorder the augmented batch according to the chosen sorting strategy to maintain genomic context:

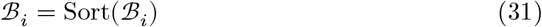

where the sorting strategy preserves the local genomic context as described in the input representation section.
6. **Final prediction:** After *n* refinement steps, a final forward pass is made using the target locations to generate the final predictions:

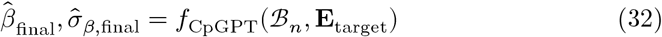

This approach leverages the model’s uncertainty estimates to selectively incorporate its most confident predictions into subsequent iterations. Crucially, it maintains a fixed step size *k* across iterations while adapting the initial candidate pool size 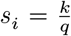 to ensure that after filtering, approximately *k* sites are added at each step. The uncertainty quantile threshold *q* determines the strictness of the filtering process; smaller values (e.g., 0.05) result in more selective filtering and require larger initial candidate pools, while larger values (e.g., 0.25) are less selective but require sampling fewer initial candidates.

Our experimental results show that this iterative refinement process leads to progressive improvements in prediction accuracy, with diminishing returns and even detrimental effects after the accumulated predictions reach approximately the same size as the original input; this likely occurs because accumulated prediction noise begins to outweigh the benefit of additional context [43].

### 4.6 Mortality predictor

To develop CpGPTGrimAge3, we started with the FHS dataset for both training and validation. We calculated all GrimAge2 and CpGPT_2M_-Proteins proxies within the dataset. For feature selection, a 5-fold cross-validation with a greedy search was utilized. Beginning with age plus a single proxy from all available proxies, proxies were added one by one until the validation z-score, corrected by age, no longer improved. This process resulted in the selection of seven GrimAge2 proxies and 23 CpGPT_2M_-Proteins proxies. These proxies, along with age, compose the input features. Finally, for the final prediction in unit of time, the Cox risk score is converted back to a year scale to align with the mean and standard deviation of the training cohort.

### 4.7 Statistics

All p-values two-sided unless otherwise stated. Any covariate used for any test are explicitly mentioned. All standard assumptions for the statistical apply unless stated otherwise.

#### 4.7.1 Clustering metrics

To quantitatively evaluate embedding quality, we computed three complementary clustering metrics: energy distance (E-distance), Adjusted Rand Index (ARI), and Normalized Mutual Information (NMI). E-distance was computed with the *scperturb* python package; for comparisons between models, a two-sided Mann-Whitney U test was applied to obtain p-values. ARI and NMI were computed against ground-truth labels: embeddings were first reduced to 50 principal components via PCA, after which KMeans clustering was applied with *k* set to the number of unique ground-truth classes (*n*_*init* = 10, deterministic seed). ARI and NMI compared the KMeans-assigned cluster labels to the ground-truth annotations. For each metric, models were ranked within each annotation category (1 = best, 3 = worst), and overall rankings were computed as the mean rank across all metrics and categories. Implementations used *scikit-learn* `KMeans, adjusted_rand_score`, and `normalized_mutual_info_score`.

#### 4.7.2 Intraclass correlation

Intraclass correlation coefficient was calculated with the *pengouin* python package. Except for the GSE55763 replicate analysis (used only for ICC), all statistical tests used distinct participants/samples.

#### 4.7.3 UMAP

To compute the UMAP, we initially reduce the data using the first 50 principal components from a PCA. Subsequently, the UMAP is determined using 30 neighbors via the *umap* python package.

#### 4.7.4 Zero-shot reference mapping

Zero-shot reference mapping was performed by transferring tissue-type labels from a well-annotated reference dataset to an unlabeled query dataset using *k*-nearest neighbor classification in the pretrained embedding space. Both the reference (AltumAge, 13,505 samples) and query (Hannum, 656 samples) datasets were passed through the pretrained CpGPT encoder without any additional finetuning to obtain sample-embedding vectors. An exact nearest-neighbor index was built on the reference embeddings using the *FAISS* library [56] with L2 (Euclidean) distance. For each query sample, *k* = 5 nearest reference neighbors were retrieved, and the most frequent tissue-type label among those neighbors was assigned by majority vote. Prior to mapping, rare tissue categories in the reference dataset whose cumulative frequency fell below 80% of samples were collapsed into an “other” category to reduce noise from underrepresented classes. The same procedure was applied to the GSE151355 dataset (20 paired blood and brain samples from Parkinson’s disease patients, 450k array), which was held out from all CpGCorpus splits and absent from the AltumAge reference, serving as an independent validation of cross-tissue classification accuracy.

#### 4.7.5 Mortality association analysis

To evaluate the association between CpGPTGrimAge3 and all-cause mortality, we trained the model in the Framingham Heart Study (FHS) cohort and evaluated performance in three independent cohorts: Massachusetts General Brigham (MGB), Women’s Health Initiative (WHI), and the Baltimore Longitudinal Study of Aging (BLSA). For each cohort, Cox proportional hazards regression models were fit with mortality as the outcome and chronological age included as a covariate. Hazard ratios (HRs) were calculated per standard deviation increase in clock score to enable direct comparison across clocks. Cox proportional hazards analyses were performed using the coxph() function from the *survival* package (version 3.6.4) in R (version 4.4.0).

#### 4.7.6 Morbidity association analysis

To evaluate associations between CpGPTGrimAge3 and age-related disease outcomes, we performed both cross-sectional and longitudinal analyses in independent cohorts. For prevalent disease analyses, logistic regression models were fit using disease status as the outcome and standardized clock score as the predictor, adjusting for chronological age. In MGB, analyses were conducted across nine disease categories: cancer, chronic obstructive pulmonary disease (COPD), cardiovascular disease (CVD), congestive heart failure (CHF), coronary artery disease (CAD), stroke, type 2 diabetes (T2D), chronic kidney disease (CKD), and cognitive decline. In BLSA, analyses were conducted across six disease categories: hypertension, myocardial infarction (MI), angina, stroke, T2D, and cancer. Odds ratios (ORs) per standard deviation increase in clock score were reported.

For longitudinal disease prediction, Cox proportional hazards models adjusting for chronological age were used to assess associations between clock scores and time-to-incident disease in MGB. In BLSA, logistic regression models were used to evaluate associations between baseline clock scores and disease incidence at two-year follow-up. Hazard ratios (HRs) and odds ratios (ORs) were calculated per standard deviation increase in clock score. For visualization, effect sizes were scaled within each disease outcome and displayed as heatmaps, enabling direct comparison of relative performance across clocks and disease endpoints. All statistical analyses were performed in R (version 4.4.0).

### 4.8 Baselines

#### 4.8.1 Statistical imputation baselines

Four non-learned baselines were used for methylation imputation benchmarking. The *sample mean* baseline imputes each missing CpG with the per-sample mean of all observed beta values in that sample. The *sample chromosome mean* extends this by computing the mean of observed sites per chromosome within each sample, so that missing CpGs are imputed with the chromosome-specific mean of the same sample. The *population chromosome mean* uses per-chromosome mean beta values computed across all training samples in CpGCorpus, independent of the test sample. Training statistics were precomputed from memory-mapped training shards and stored as compressed arrays. The *population locus mean* is the most informative population baseline, using the per-CpG site mean beta value across all training samples, keyed by chromosome index and genomic position; it is limited to CpG sites observed during training so it was not used for tasks in which a CpG site was completely unseen and not part of training (e.g. EPIC to mammalian conversion, 450k to MSA conversion).

#### 4.8.2 mLiftOver

For direct comparison on the EPIC-to-Mammalian array conversion task, we used mLiftOver [38] with pre-trained elastic net coefficients fitted on 141 paired human blood samples to convert from the EPIC to the mammalian array (`fit_EPICtoMM_v1_N94_N=141.rds`). Missing EPIC probes were filled with *β* = 0.5 (corresponding to *M* = 0).

#### 4.8.3 DeepCpG

For single-cell methylation imputation comparison on the sciMETv3 dataset, we reimplemented DeepCpG [41] in PyTorch. The architecture follows the original publication: a DNA module (multi-scale CNN over 1,001 bp windows with filter sizes [128, 256] and kernel sizes [11, 7]), a CpG module (bidirectional GRU with 128 hidden units per direction over 25 neighboring cells), and a joint module (256 hidden units, dropout 0.2). The model was trained using PyTorch Lightning with a batch size of 256, learning rate of 1 × 10^−3^, maximum 50 epochs with early stopping (patience = 5 on validation loss), and mixed precision (16-bit). The best checkpoint was selected by validation loss. The reference genome used was GRCh38 (Ensembl release 113). For fair comparison, CpGPT’s held-out masking scheme was applied to DeepCpG so that both models predicted identical sets of held-out CpG sites.

#### 4.8.4 ElasticNet, logistic regression and MLP baselines

The ElasticNet, logistic regression and MLP baselines were calculated with *scikit-learn* version 1.7.0 using the classes `ElasticNet, LogisticRegression`, and `MLPRegressor` or `MLPClassifier` with default parameters respectively.

## 5 Code and data availability

CpGCorpus is publicly available for access in the S3 bucket s3:// cpgpt-lucascamillo-public/data/cpgcorpus/raw. Model weights and dependencies are accessible via HuggingFace https://huggingface.co/lucascamillomd/cpgpt-models.

The code and the tutorials are available in our GitHub repository under http://github.com/lucascamillomd/CpGPT. CpGPTGrimAge3 is available on pyaging http://github.com/lucascamillomd/pyaging.

The datasets used for the analyses described in this manuscript were obtained from dbGaP at http://www.ncbi.nlm.nih.gov/sites/entrez?db=gap through dbGaP accessions phs000200 and phs000007. BLSA data is available upon reasonable request to the Intramural Research Program of the National Institute on Aging. Access applications can be submitted at https://www.blsa.nih.gov/.

## 6 Acknowledgements

We would like to thank the Zhou lab for their comprehensive, publicly available annotation of the Illumina methylation arrays [52]. We would also like to thank Alexander Mathiasen for the fruitful conversations. Moreover, part of this work was supported by the Gruber Science Fellowship at Yale University (R.S.), the National Institute on Aging (A.H.C.), and an AI at Yale Seed Grant (A.H.C.).

The WHI program is funded by the National Heart, Lung, and Blood Institute, National Institutes of Health, U.S. Department of Health and Human Services through contracts HHSN268201600018C, HHSN268201600001C, HHSN268201600002C, HHSN268201600003C, and HHSN268201600004C. This manuscript was not prepared in collaboration with investigators of the WHI, has not been reviewed and/or approved by the Women’s Health Initiative (WHI), and does not necessarily reflect the opinions of the WHI investigators or the NHLBI. The Framingham Heart Study is conducted and supported by the National Heart, Lung, and Blood Institute (NHLBI) in collaboration with Boston University (Contract No. N01-HC-25195, HHSN268201500001I and 75N92019D00031). This manuscript was not prepared in collaboration with investigators of the Framingham Heart Study and does not necessarily reflect the opinions or views of the Framingham Heart Study, Boston University, or NHLBI. The DNA methylation data provided in this data set were supported by funding from the NIH including funds from the NHLBI, Division of Intramural Research (D. Levy, PI) and the NIH Director’s Challenge Award (D. Levy, PI). The BLSA study is conducted by the Intramural Research Program of the NIA, part of the NIH at the US Department of Health and Human Services. The authors also would like to thank the staff and participants of the BLSA.

## 7 Conflicts of interest

L.P.D.L.C. has share options at Shift Bioscience Ltd and has received consulting fees from TruDiagnostic. R.S. has received consulting fees from TruDiagnostic, LongevityTech.fund and Cambrian BioPharma. A.H.C. has received consulting fees from TruDiagnostic and FOXO Biosciences. B.W. serves as a scientific advisor to Shift Bioscience, Vevo Therapeutics, Deep Genomics. B.W. receives consulting fees from Arsenal Bioscience, Viecure Inc. S.H. is a founder and paid consultant of the non-profit Epigenetic Clock Development Foundation, which licenses these patents. S.H. was a Principal Investigator at Altos Labs until March 2026 and holds equity in Altos Labs, Inc. The Regents of the University of California are the sole owner of patents and patent applications directed at epigenetic biomarkers and the mammalian methylation array platform for which S.H. is a named inventor.

## Supplementary Table Legends

All supplementary tables are provided in the accompanying Excel file (`Supplementary_Tables.xlsx`), with each table on a separate tab.

**Supplementary Table 1**. DNA language model and context length ablation. Test beta MAE and test loss on the CpGCorpus held-out set for six DNA sequence encoders across three flanking context window lengths. All models use the CpGPT_2M_ architecture trained for 100,000 steps with chromosomal sorting.

**Supplementary Table 2**. Architecture ablation. Test beta MAE and test loss for three configurations of the CpGPT_2M_ model using Nucleotide Transformer V2 500M with context length of 2001 bps, trained for 100,000 steps.

**Supplementary Table 3**. Training duration scaling. Test and validation beta MAE as a function of the number of training steps for the reference CpGPT_2M_ configuration (Nucleotide Transformer V2 500M, context length of 2001 bps, with transformer and chromosomal sorting).

**Supplementary Table 4**. Quantitative evaluation of locus embedding quality across six genomic annotation categories. For each category, embeddings from CpGPT_2M_, CpGPT_100M_, and NTv2_500M_ are compared using three metrics: Adjusted Rand Index (ARI), Normalized Mutual Information (NMI), and median Energy distance (E-dist). Models are ranked per category (1 = best discrimination). CpGPT_2M_ achieves the best overall average rank (1.22), followed by CpGPT_100M_ (2.28) and NTv2_500M_ (2.50).

**Supplementary Table 5**. Clustering metrics summary across datasets and methods. Adjusted Rand Index (ARI) and Normalized Mutual Information (NMI) for CpGPT_2M_, CpGPT_100M_, fine-tuned variants, and baseline methods across locus-level (6 genomic annotation categories) and sample-level (AltumAge, human RRBS atlas, mammalian, sciMETv3) datasets.

**Supplementary Table 6**. Zero-shot reference mapping of OSKM reprogramming samples (GSE54848) to the CpGCorpus. Each sample from the fibroblast-to-iPSC reprogramming time course was mapped to its nearest neighbor in the CpGCorpus embedding space using CpGPT_100M_. Day 0 samples (pre-induction fibroblasts) map to somatic primary human dermal fibroblast (HDF) entries, while day 28 samples map to induced pluripotent stem cell (iPSC) entries. Day 15 is the last time point at which samples retain a fibroblast-associated mapping, consistent with previous reports of identity loss during reprogramming [32].

**Supplementary Table 7**. CpG probe universe partitioned by chromosome. The training set covers 1,174,502 of the 1,186,366 probes in the combined Illumina manifest (99.0%). The remaining 11,864 probes (1.0%) are held out and only appear in the test split, allowing evaluation of the model’s ability to generalize to unseen genomic positions.

**Supplementary Table 8**. Held-out CpG probes. Full list of the 11,864 probe IDs reserved for the test split (companion to Supplementary Table 7).

**Supplementary Table 9**. Tissue-stratified imputation performance on the CpGCorpus test set. Mean absolute error (MAE) is reported per tissue category for CpGPT_2M_, CpGPT_100M_, and statistical baselines (sample mean, sample chromosome mean, population chromosome mean, population locus mean^†^) across three evaluation regimes: seen → seen, seen → partially unseen, and seen → fully unseen. CpGPT consistently outperforms statistical baselines across all tissues. Performance is lowest for Fibroblasts/Skin, likely reflecting their underrepresentation in the blood-dominated training set. ^†^ Population locus mean is undefined for CpGs absent from the training set, so this baseline is reported as NA for the partially-unseen and fully-unseen splits.

**Supplementary Table 10**. Chain-of-thought iterative refinement results on the Hannum dataset. For both CpGPT_2M_ and CpGPT_2M_-Hannum, beta-value MAE is reported as a function of the number of refinement steps (0–many), step size (500, 1,000 or 2,000), and uncertainty quantile (5th, 10th, 25th, 50th and 100th percentile). Starting from 5,000 MSA CpG sites, CpGPT_2M_ reaches its lowest MAE (0.0533 ± 0.0023) after 9 refinement steps of size 500 under the 5th-percentile filter; the finetuned CpGPT_2M_-Hannum reaches 0.0410 ± 0.0027 after 6 steps under the same filter.

**Supplementary Table 11**. Per-dataset chronological-age prediction performance on the AltumAge multi-tissue test set. For each of the 29 test studies (with sample size and tissue annotation), MAE, MSE and Pearson correlation are reported for CpGPT_2M_-Age, ElasticNet, and an MLP baseline. CpGPT_2M_-Age achieves the lowest MAE on the majority of studies, demonstrating that the headline correlation (Pearson *r* = 0.9847 on Figure 4a) reflects consistent generalization across tissues, age ranges and array platforms rather than a few easy datasets. Pearson *r* is left blank for datasets in which all samples share the same chronological age (zero variance in the ground truth), since the correlation coefficient is mathematically undefined in that case; MAE and MSE remain well defined and are reported.

**Supplementary Table 12**. Per-protein prediction performance for all 322 plasma proteins in the Biology of Aging (BoA) challenge dataset. CpGPT achieves statistically significant predictions (*p* < 0.05) for 148 of 322 proteins (46%), compared to 120 (37%) for ElasticNet. The best-predicted proteins are enriched for neurodegeneration-associated markers (NEFL, NEFH, GFAP, TREM2, pTau-181, MAPT), immune signalling molecules (CCL11, CD40, TNFRSF1A/1B, IL15RA), and extracellular matrix regulators (IGFBP7, MMP3, CHI3L1).

**Supplementary Table 13**. Dataset splits. A detailed list of GEO Series (GSE) entries included in each train, validation, and test split, along with the number of samples and CpG sites per entry.

**Supplementary Table 14**. All-cause mortality associations across cohorts. Hazard ratios (HRs) per standard deviation increase in clock score, with 95% confidence intervals (lower and upper limits), from age-adjusted Cox proportional hazards models for CpGPT-GrimAge3 and eight established epigenetic clocks (PCHorvath1, PCHannum, PhenoAge, PCPhenoAge, DunedinPACE, SystemsAge, GrimAgeV2, and PCGrimAge). CpGPTGrimAge3 was trained in the Framingham Heart Study (FHS) and evaluated in three independent held-out cohorts: Massachusetts General Brigham (MGB), the Women’s Health Initiative (WHI), and the Baltimore Longitudinal Study of Aging (BLSA). CpGPTGrimAge3 has the largest HR among all clocks in each of the three held-out cohorts and the second-largest in FHS.

**Supplementary Table 15**. Disease associations across cohorts, analyses, and clocks. Effect sizes between clock scores and age-related disease are reported by cohort and analysis type: cross-sectional associations with prevalent disease at baseline (odds ratios from age-adjusted logistic regression) in MGB and BLSA, longitudinal associations with time to incident disease (hazard ratios from age-adjusted Cox models) in MGB, and disease incidence at two-year follow-up in BLSA. Disease categories are cancer, COPD, CVD, CHF, CAD, stroke, type 2 diabetes (T2D), CKD, and cognitive decline in MGB, and hypertension, myocardial infarction, angina, stroke, T2D, and cancer in BLSA. Blank cells indicate categories with insufficient events for stable estimation. Across the 34 mortality and disease comparisons, CpGPTGrimAge3 ranks among the top two clocks in 30 (88.2%) and first in 18 (52.9%).

## Supplementary Figures

**Supplementary Figure 1:**
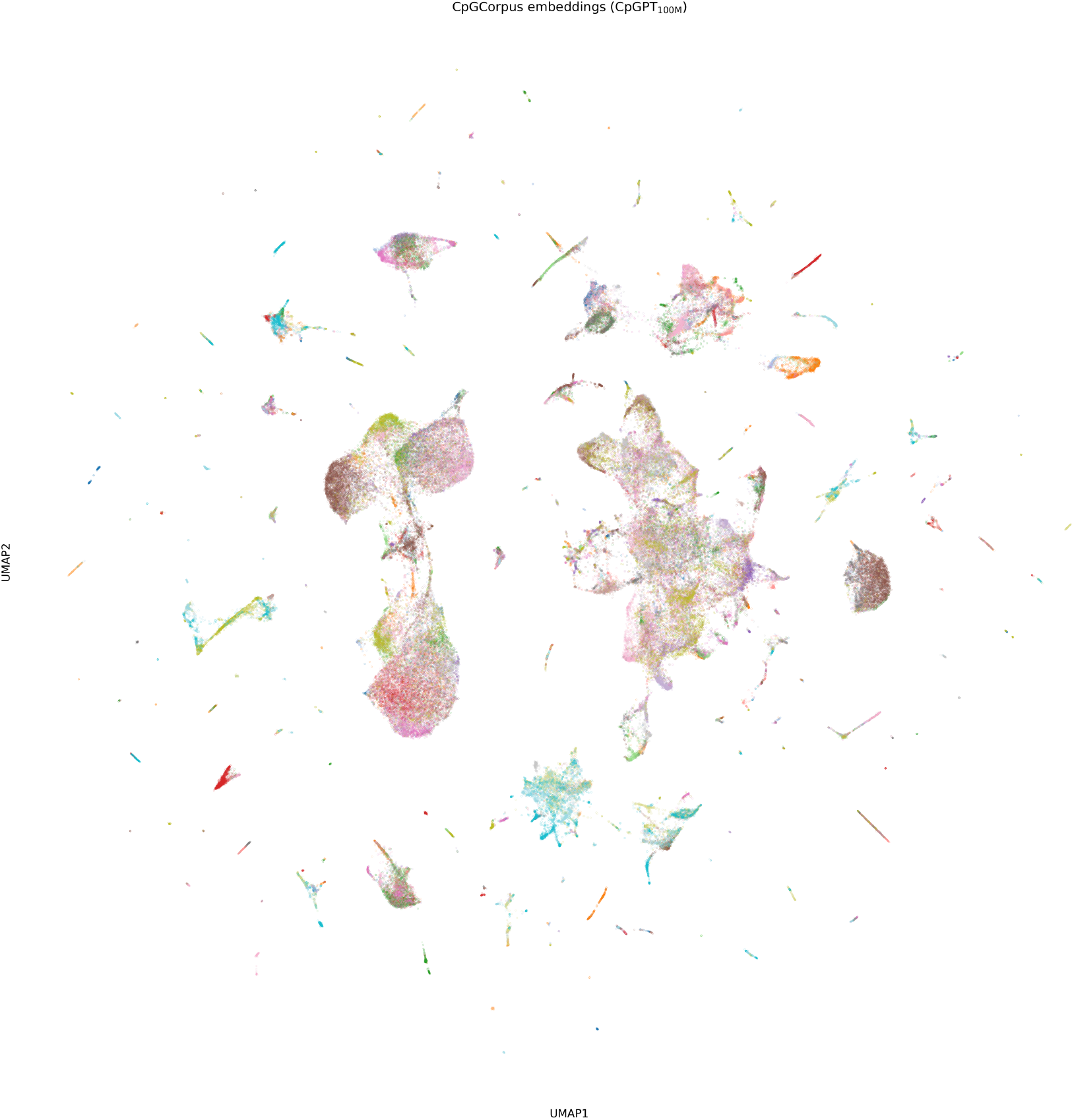
UMAP of the CpGPT_100M_ embeddings of the training set of CpGCorpus colored by dataset.

**Supplementary Figure 2:**
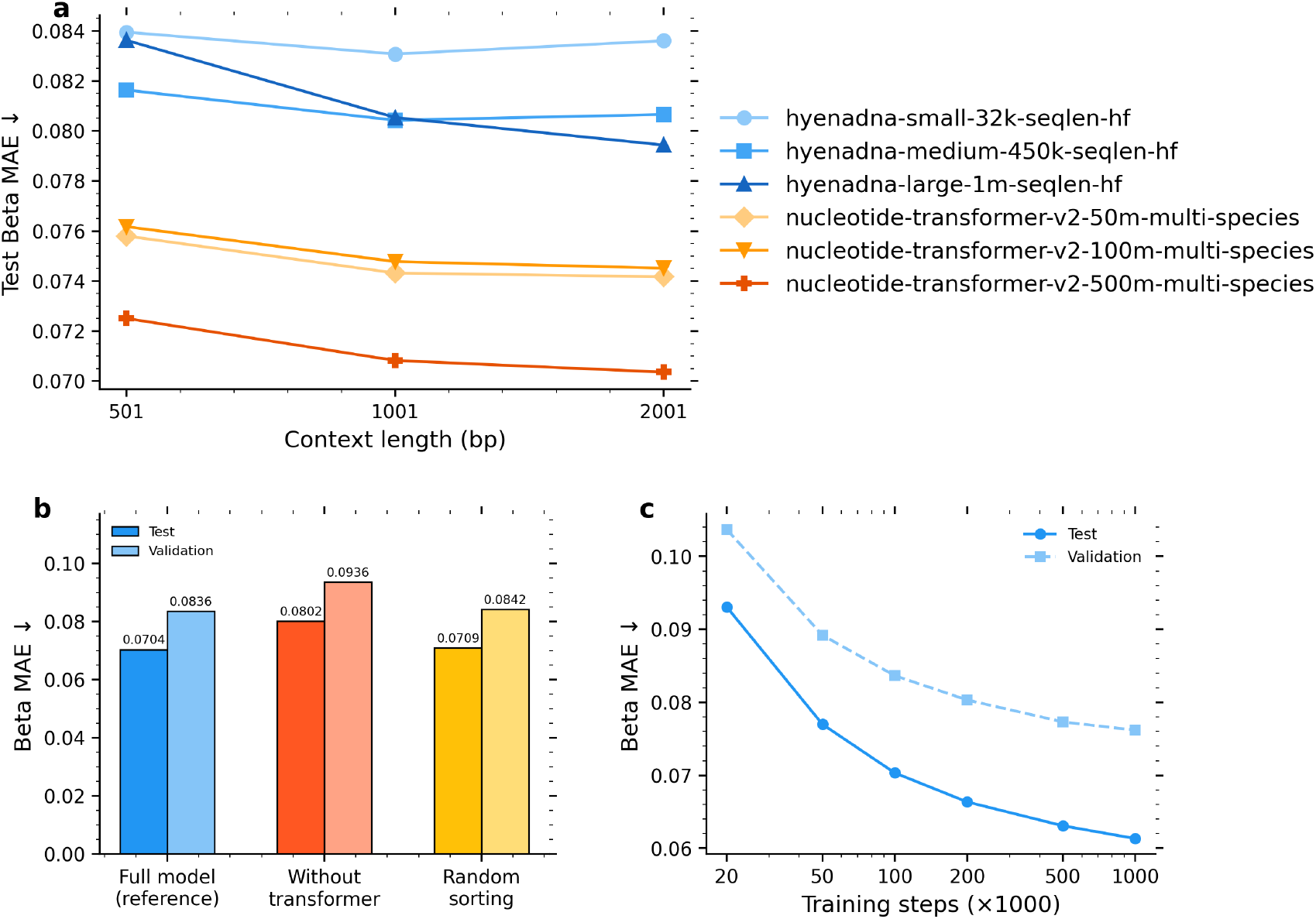
Ablation studies for CpGPT_2M_ architectural and training choices. (a) Test beta MAE on the CpGCorpus held-out set as a function of DNA language model backbone (hyenadna-small-32k-seqlen-hf, hyenadna-medium-450k-seqlen-hf, hyenadna-large-1m-seqlen-hf, nucleotide-transformer-v2-50m-multi-species, nucleotide-transformer-v2-100m-multi-species, nucleotide-transformer-v2-500m-multi-species) and flanking context window length (501, 1001, and 2001 bps). All models were trained for 100k gradient steps with chromosomal sorting enabled. (b) Architecture ablation comparing the reference configuration (nucleotide-transformer-v2-500m-multi-species, context length = 2001 bp, 100k steps) against a variant without transformer blocks and a variant with random CpG ordering instead of chromosomal sorting. Lighter bars show corresponding validation set performance. (c) Training duration scaling from 20k to 1M gradient steps for the reference configuration (nucleotide-transformer-v2-500m-multi-species, ctx = 2001 bp). The test (solid) and validation (dashed) beta MAE both decrease following an approximate power-law trajectory. See **Supplementary Tables 1–3** for the full numerical results.

**Supplementary Figure 3:**
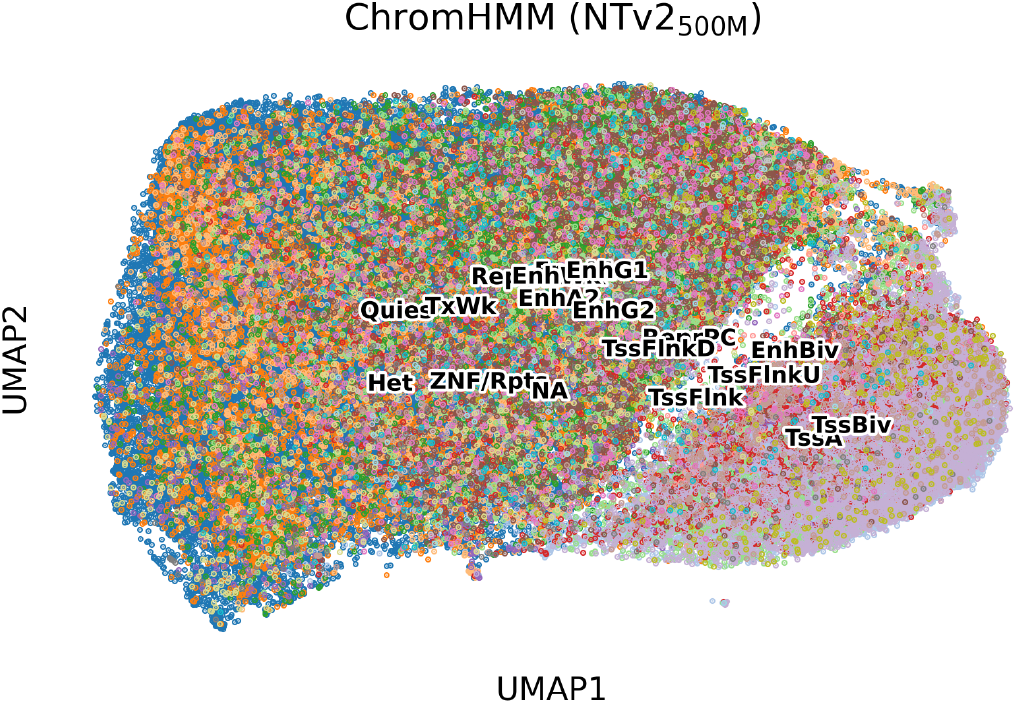
UMAP of the locus embeddings from Nucleotide Transformer V2 500M colored by ChromHMM genomic annotation

**Supplementary Figure 4:**
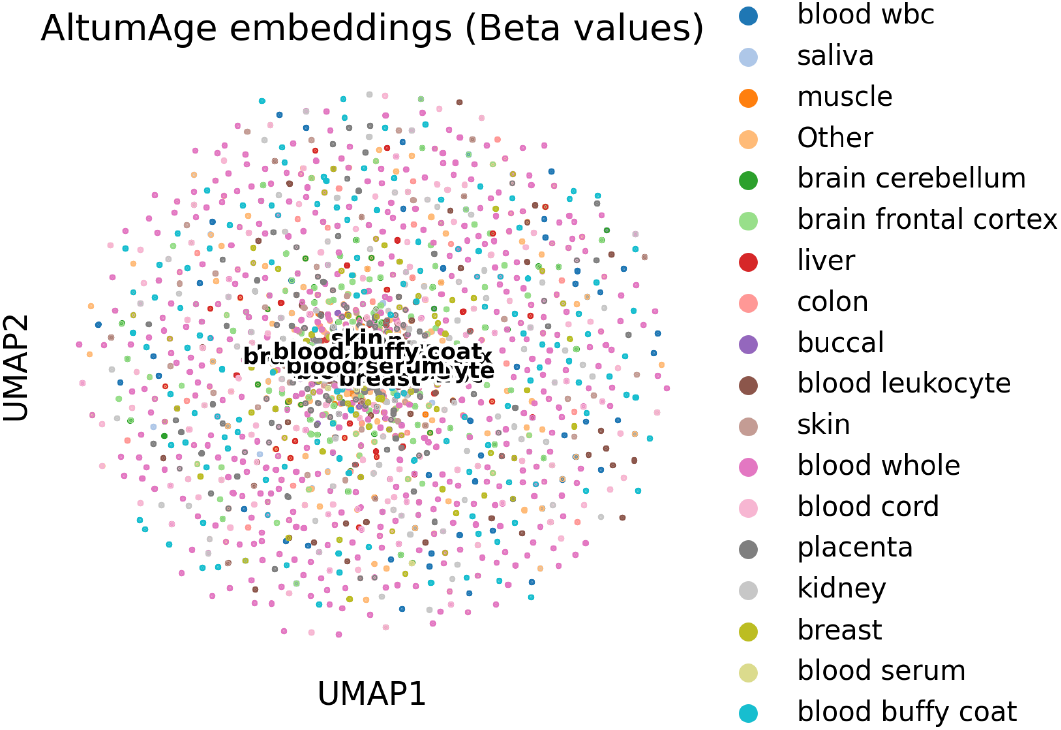
UMAP of the beta values for the AltumAge dataset colored by tissue type.

**Supplementary Figure 5:**
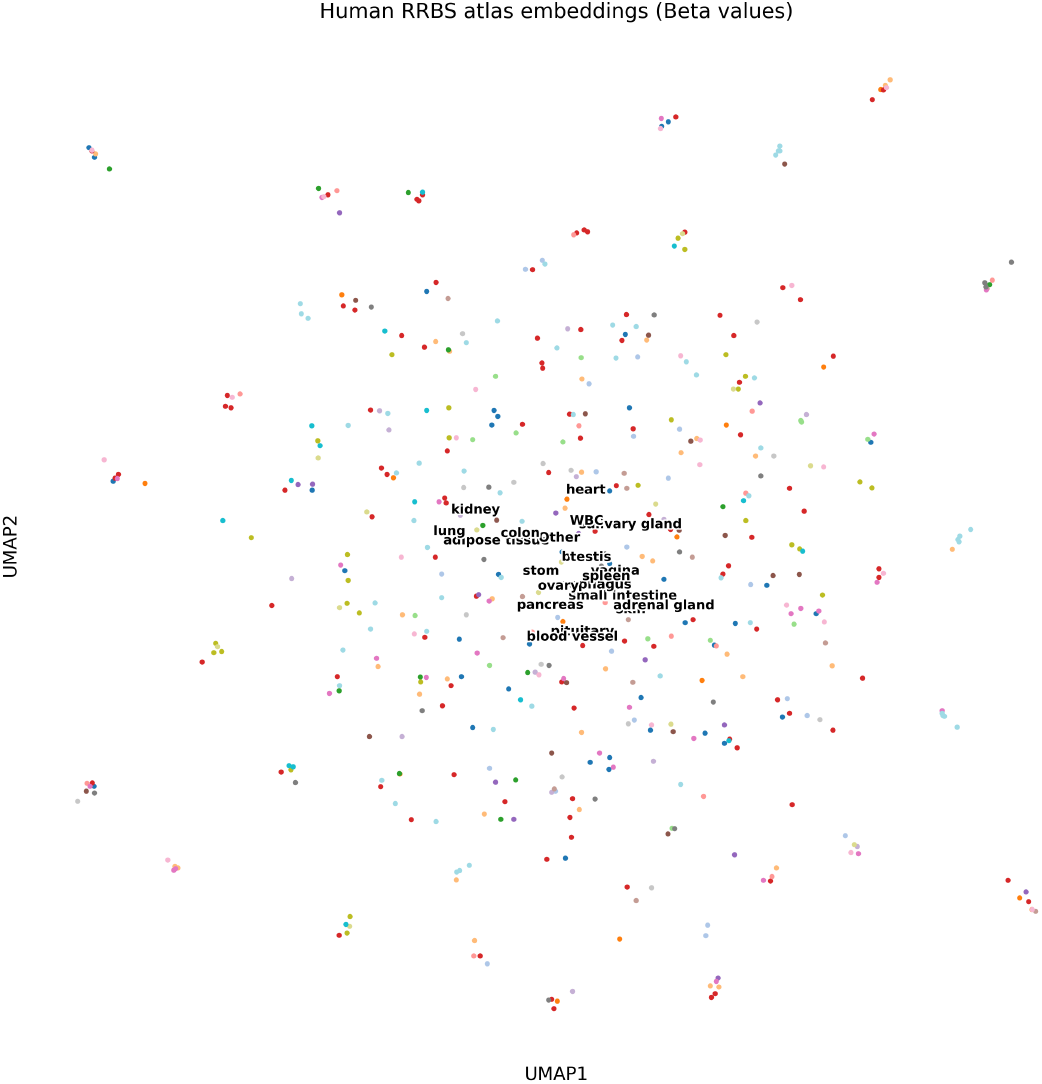
UMAP of raw beta values from the human RRBS atlas dataset (GSE233417) colored by tissue type. Raw methylation values yield weak tissue-type separation (ARI = 0.001; NMI = 0.146). The corresponding CpGPT_100M_ sample embeddings are shown in Supplementary Figure 6.

**Supplementary Figure 6:**
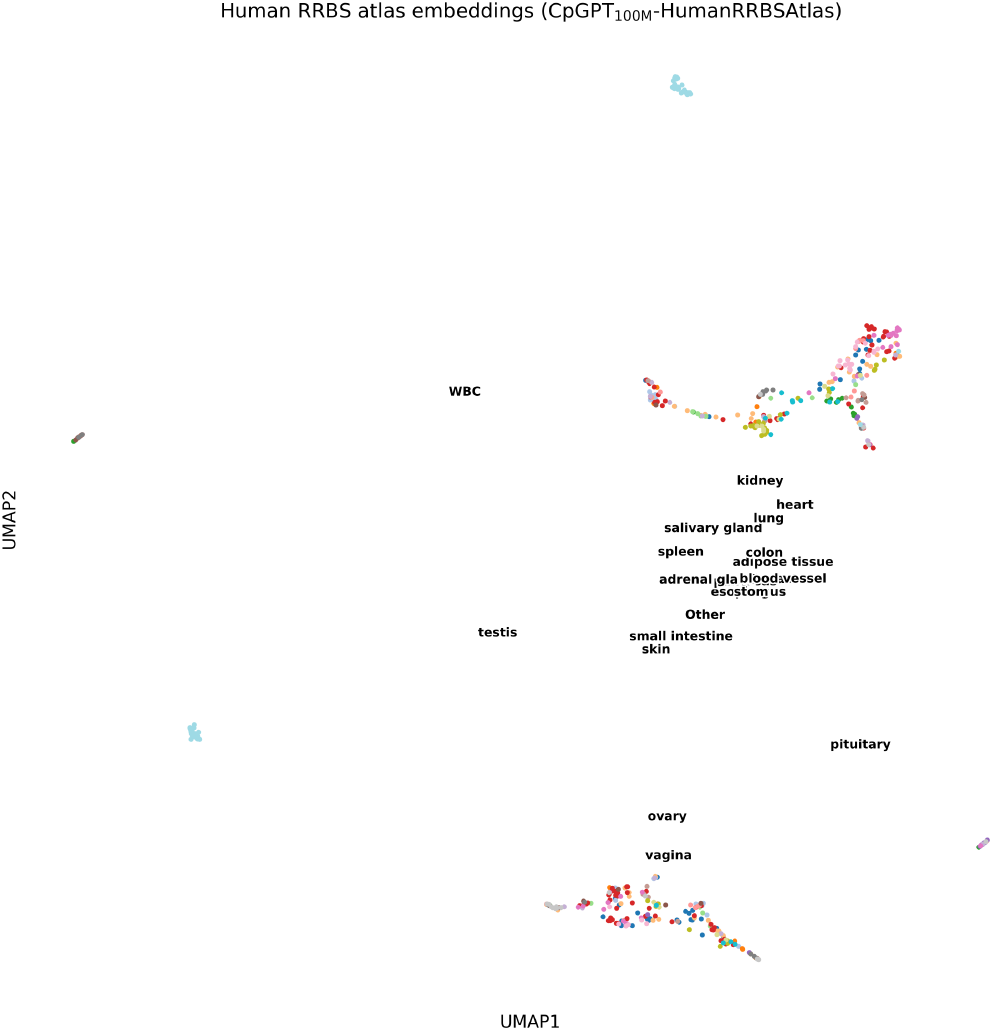
UMAP of the CpGPT_100M_ sample embeddings for the human RRBS atlas dataset (GSE233417) colored by tissue type. The CpGPT_100M_ embeddings show substantially improved tissue-type separation (ARI = 0.091; NMI = 0.332) compared to raw beta values shown in Supplementary Figure 5.

**Supplementary Figure 7:**
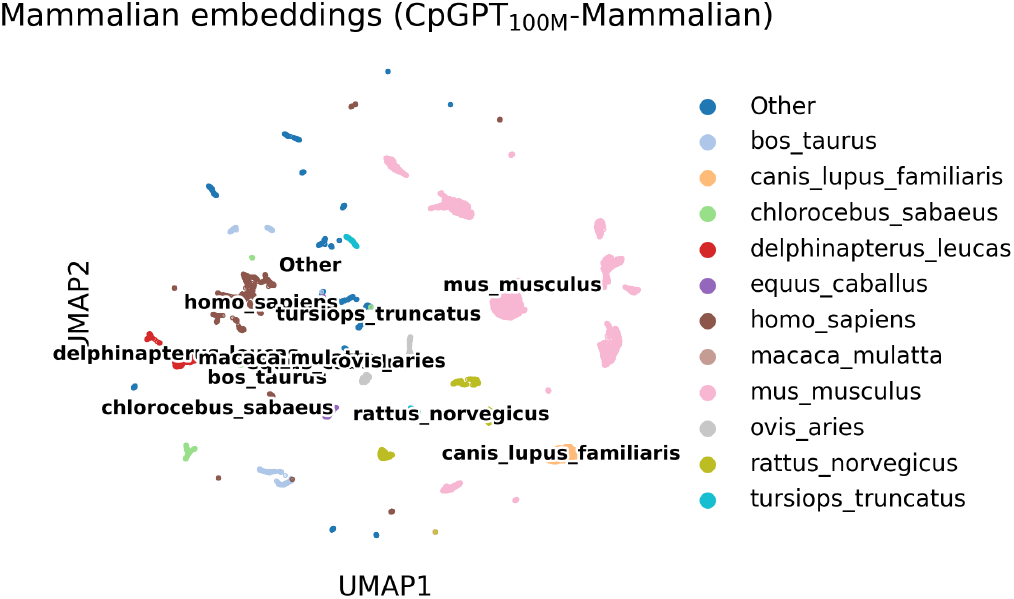
UMAP of the *fine-tuned* CpGPT_100M_-Mammalian sample embeddings for the pan-mammalian dataset (GSE223748), colored by species. In contrast to the pretrained embeddings shown in Figure 2d (where species is the dominant clustering axis), the fine-tuned model reorganizes the embedding space so that tissue type becomes the primary separator (see Section 2.2).

**Supplementary Figure 8:**
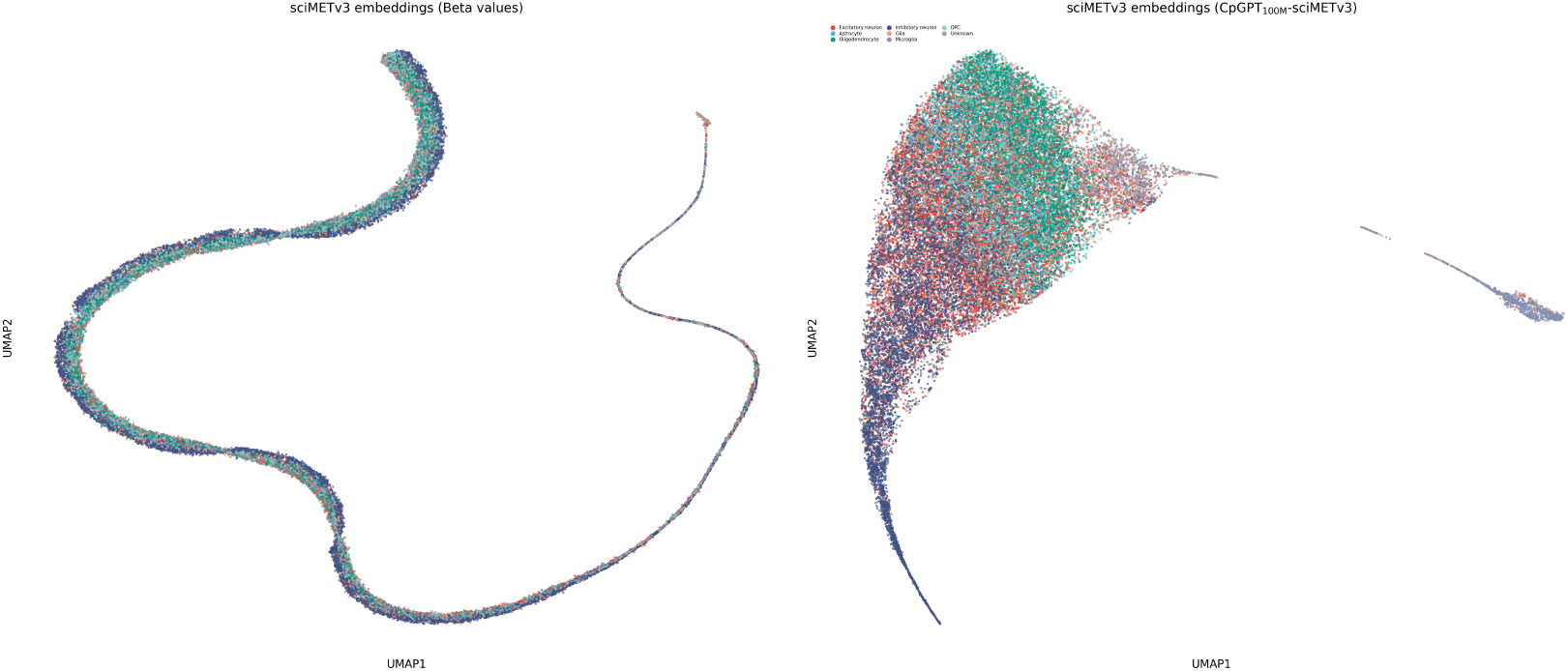
UMAP of the beta values (left) and CpGPT_100M_-sciMETv3 (right) samples embeddings of the single-cell sciMETv3 dataset.

**Supplementary Figure 9:**
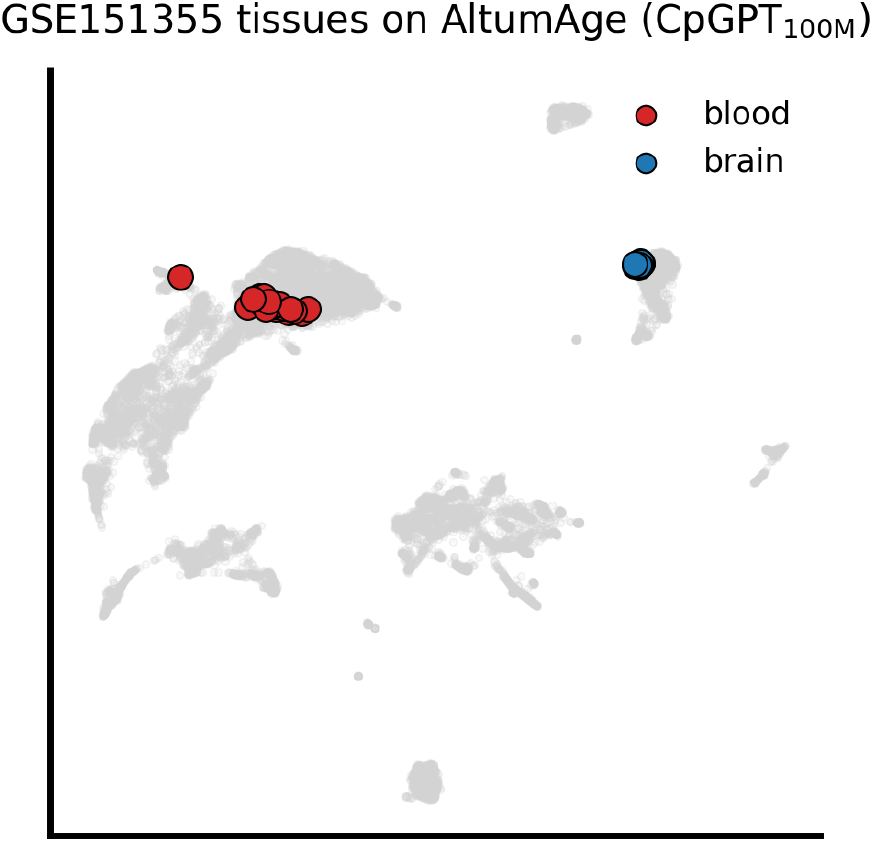
Zero-shot tissue classification of paired blood and brain samples (GSE151355) via reference mapping onto the AltumAge dataset using CpGPT_100M_ embeddings. Each point represents a sample projected into the AltumAge UMAP space (gray background). Red points: blood samples (*n* = 20); blue points: brain samples (*n* = 20). Using *k* = 5 nearest-neighbor label transfer, all 20 blood samples were classified as “blood whole” and 17 of 20 brain samples as “brain frontal cortex” (broad-match accuracy = 92.5%). This dataset was not included in CpGCorpus training, validation, or testing, nor in the AltumAge reference panel.

**Supplementary Figure 10:**
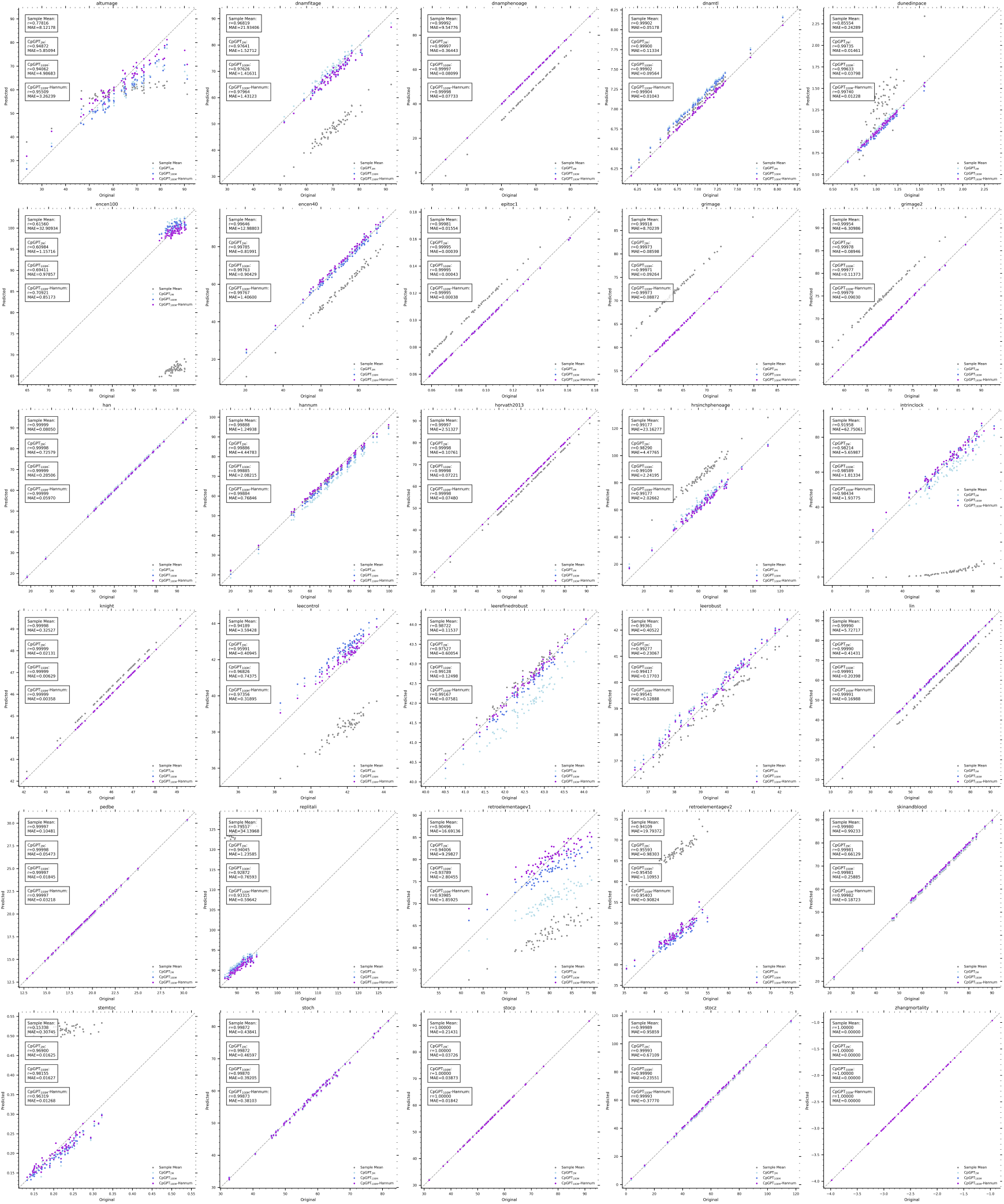
Scatterplot of 30 epigenetic clocks predicted based on different imputation methods for the 450k probes missing from the MSA array in the Hannum dataset with a mean baseline, CpGPT_2M_, CpGPT_100M_, and CpGPT_100M_-Hannum.

**Supplementary Figure 11:**
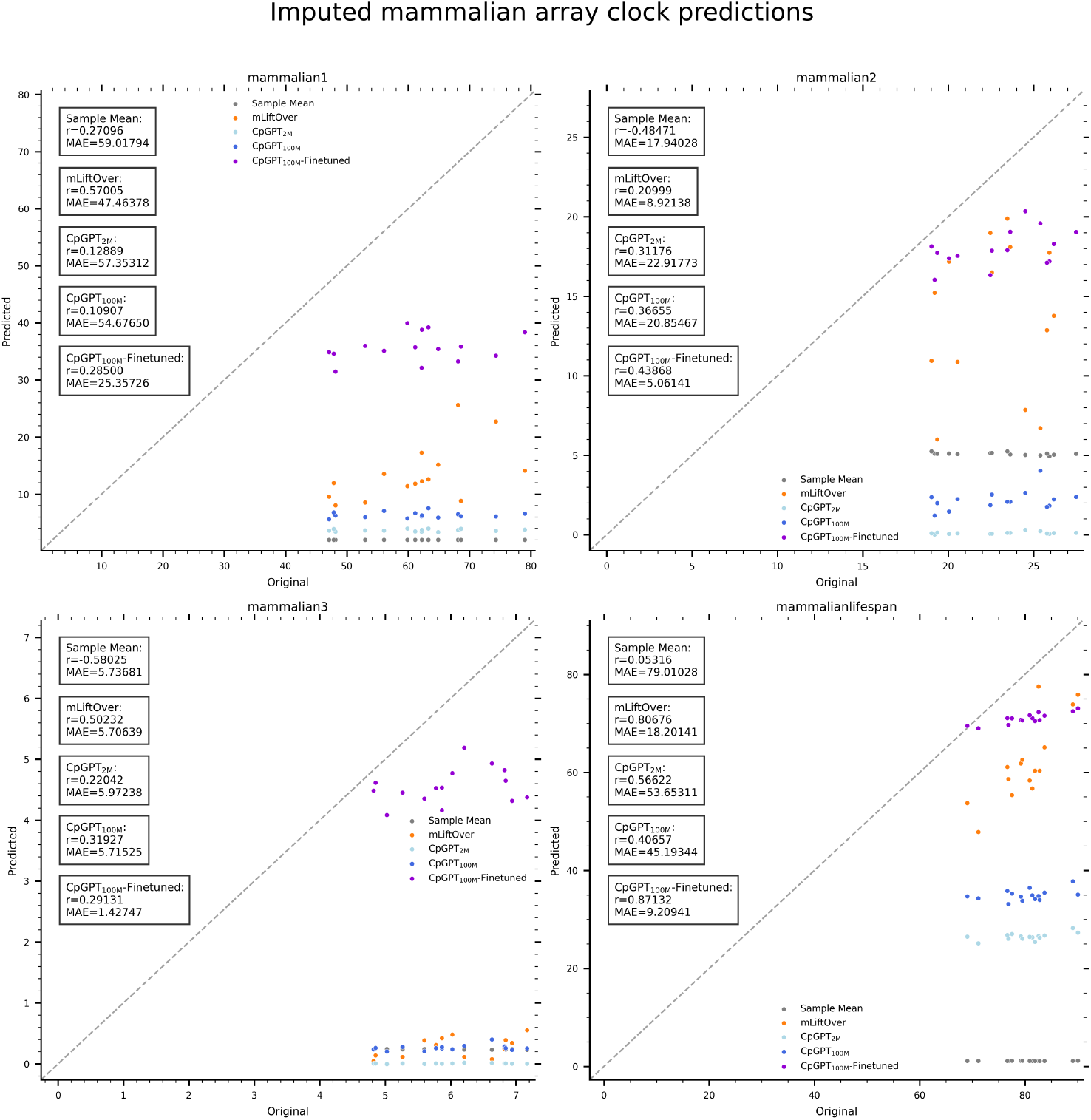
Scatterplot of four epigenetic clocks applied to the reconstructed mammalian array from EPIC data with a mean baseline, CpGPT_2M_, CpGPT_100M_, and CpGPT_100M_-EpicMammal.

**Supplementary Figure 12:**
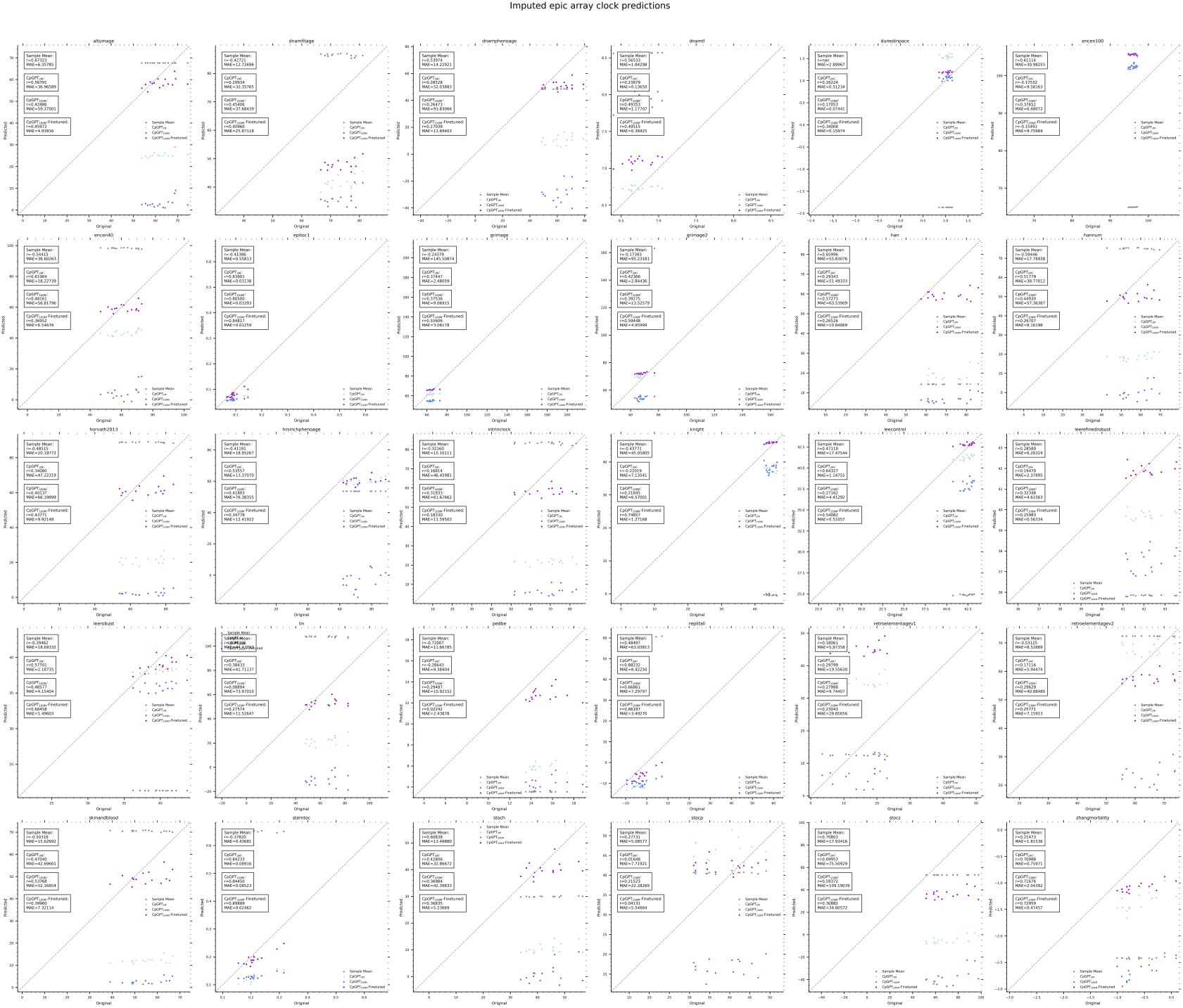
Scatterplot of 30 epigenetic clocks applied to the reconstructed EPIC array from mammalian array data with a mean baseline, CpGPT_2M_, CpGPT_100M_, and CpGPT_100M_-EpicMammal.

**Supplementary Figure 13:**
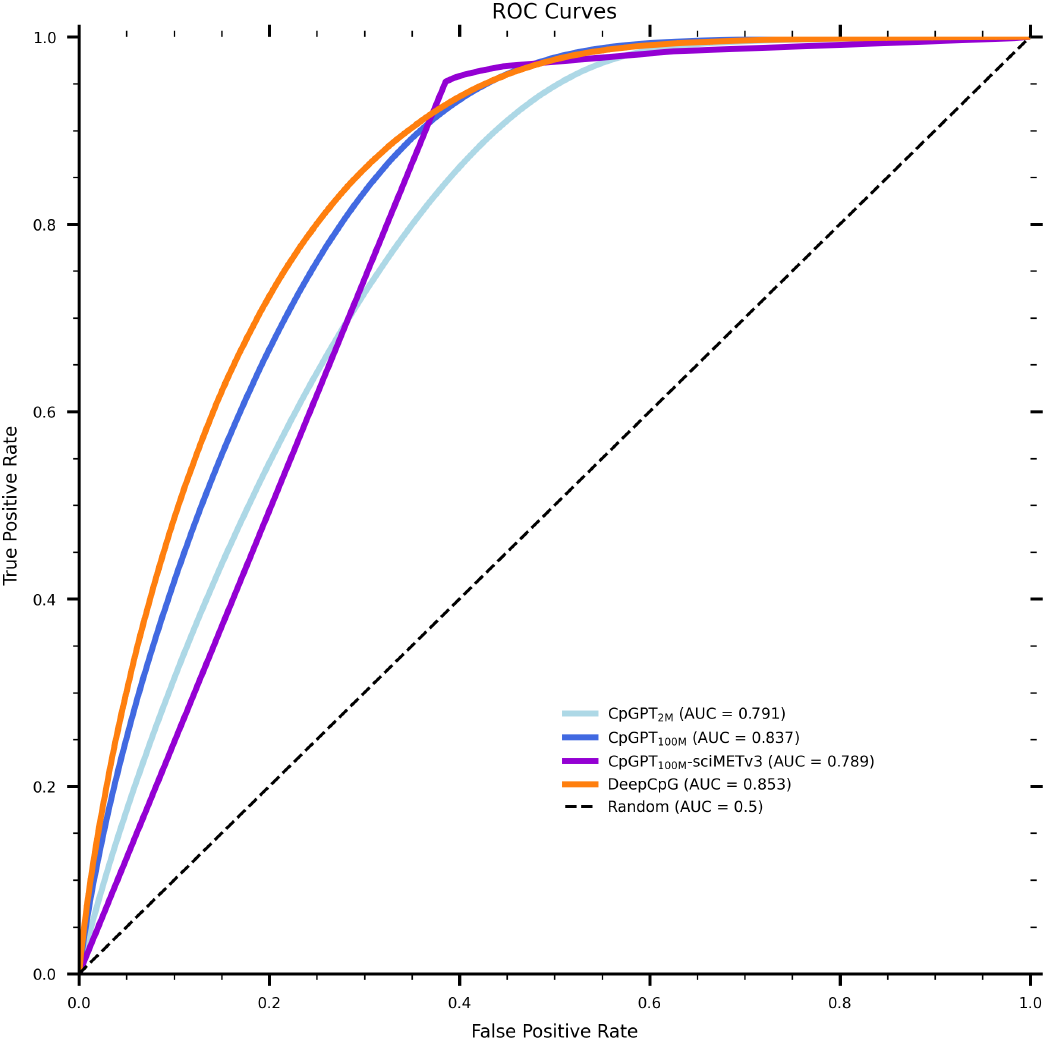
Receiver-operator characteristic (ROC) curves for single-cell methylation imputation on the sciMETv3 dataset, comparing statistical baselines (sample mean, chromosome means, locus mean), DeepCpG, CpGPT_2M_, CpGPT_100M_, and finetuned CpGPT_100M_-sciMETv3.

**Supplementary Figure 14:**
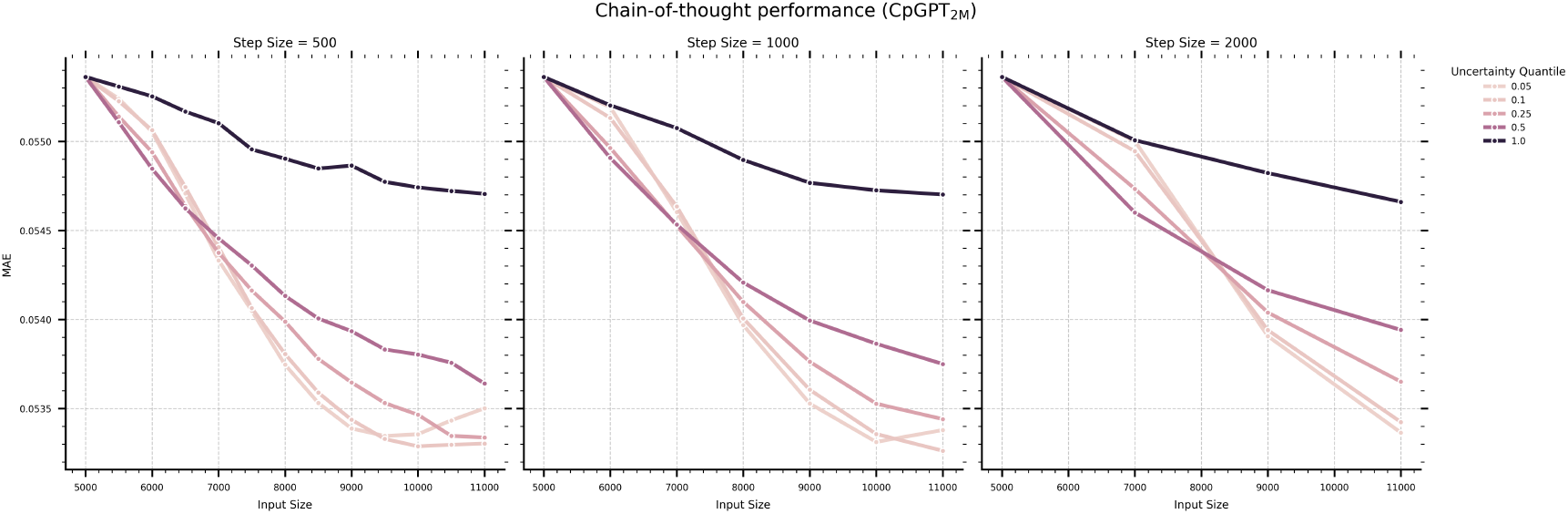
Lineplot showing the iterative refinement performance of CpGPT_2M_ in reconstructing non-MSA 450k probes in the Hannum dataset based on different step sizes and uncertainty thresholds.

**Supplementary Figure 15:**
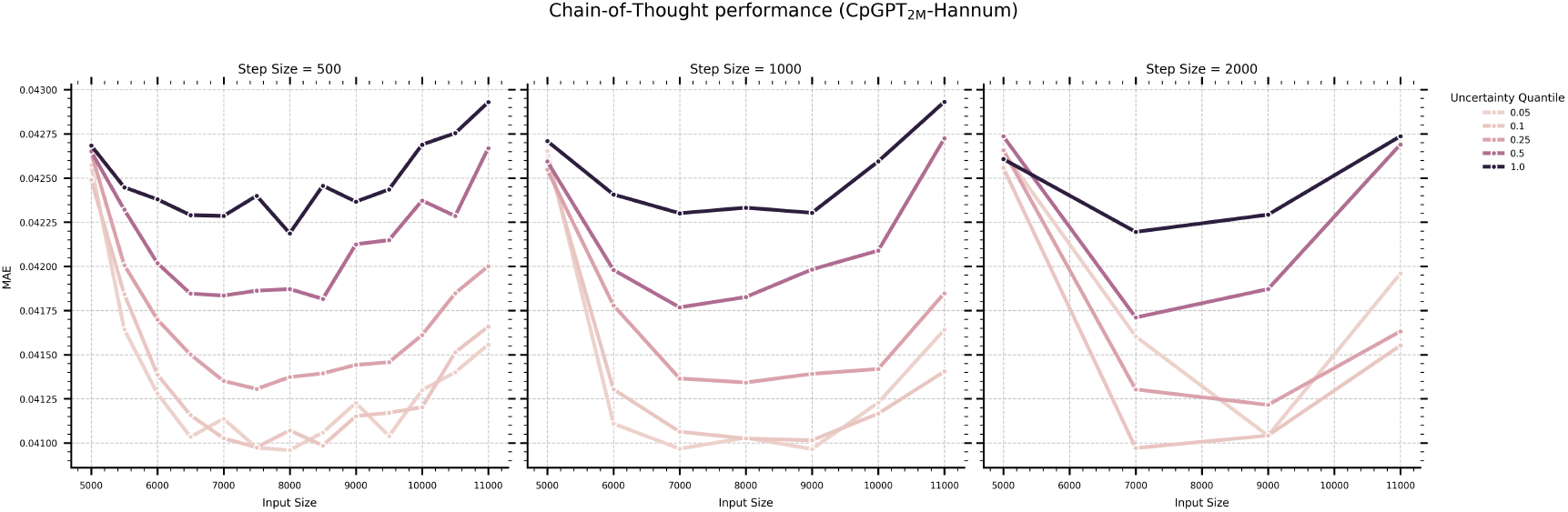
Lineplot showing the iterative refinement performance of CpGPT_2M_-Hannum in reconstructing non-MSA 450k probes in the Hannum dataset based on different step sizes and uncertainty thresholds.

**Supplementary Figure 16:**
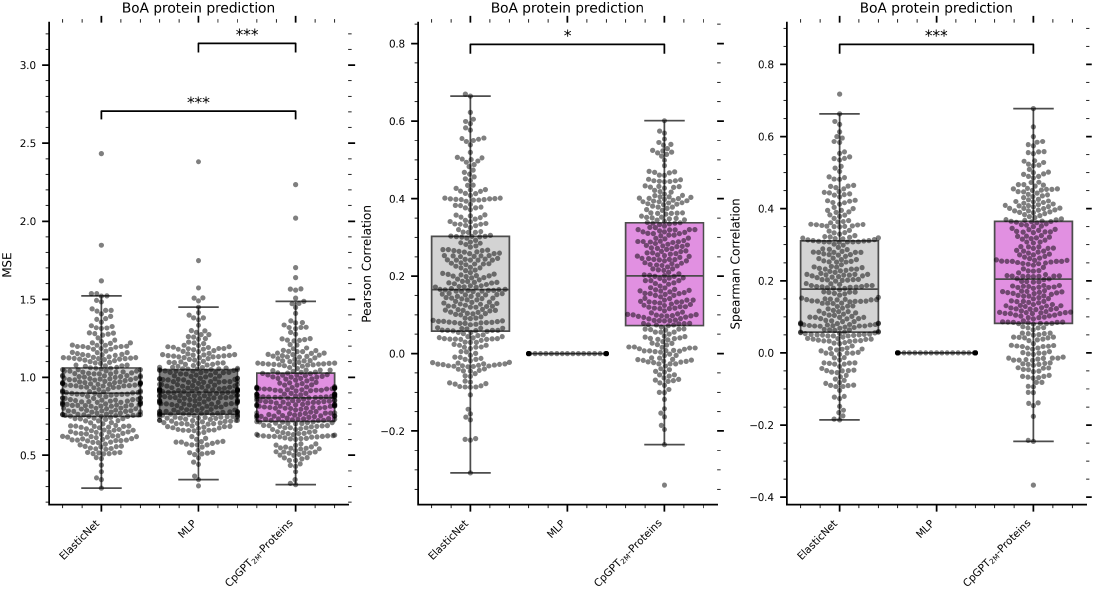
Boxplot of the mean squared error (MSE), Pearson correlation, and Spearman correlation coefficients for the prediction of 322 proteins from blood methylation of the Biomarkers of Aging Consortium dataset comparing ElasticNet, MLP, and CpGPT_2M_-Proteins.

**Supplementary Figure 17:**
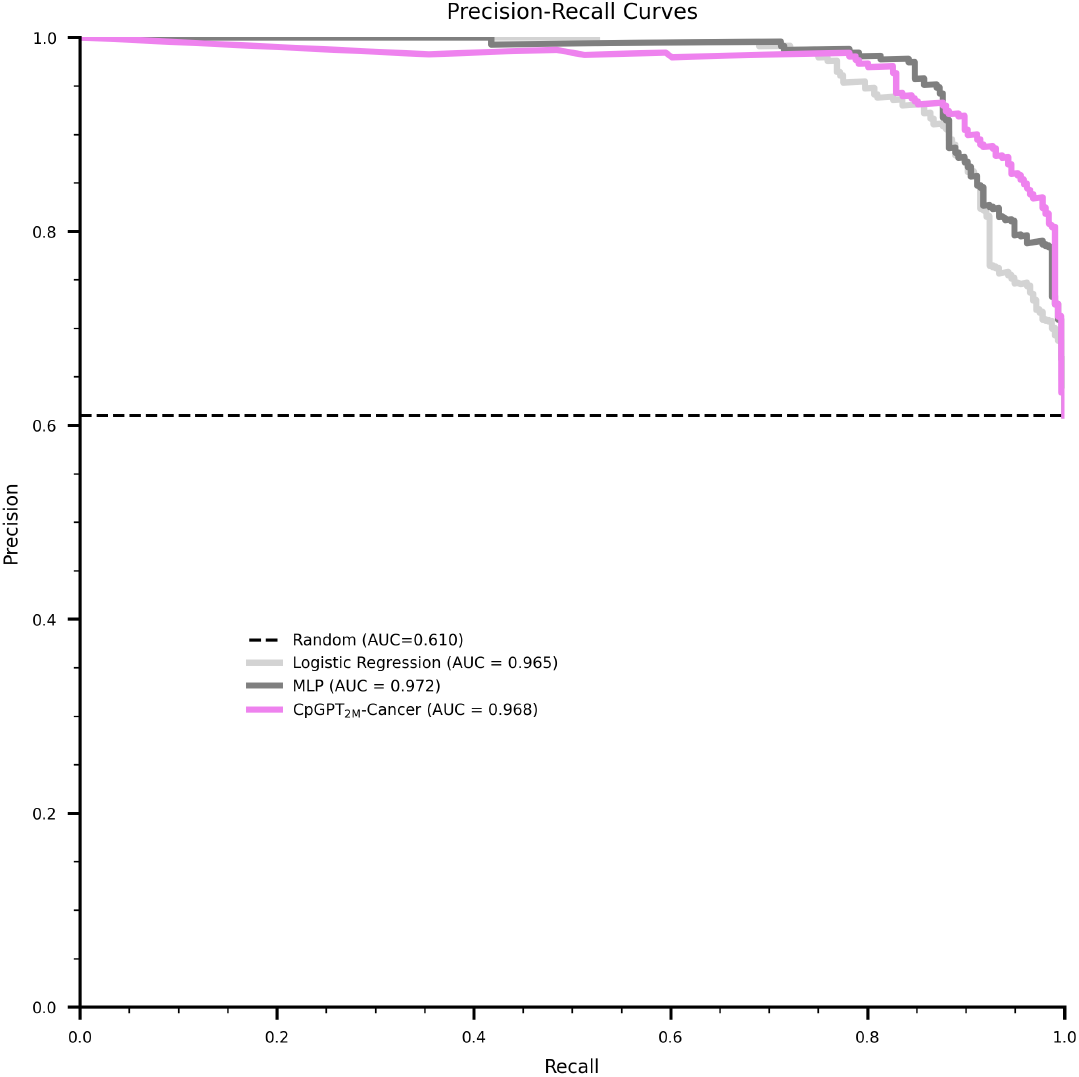
Precision-recall curve for the cancer prediction in the AltumAge dataset comparing logistic regression, MLP, and CpGPT_2M_-Cancer

**Supplementary Figure 18:**
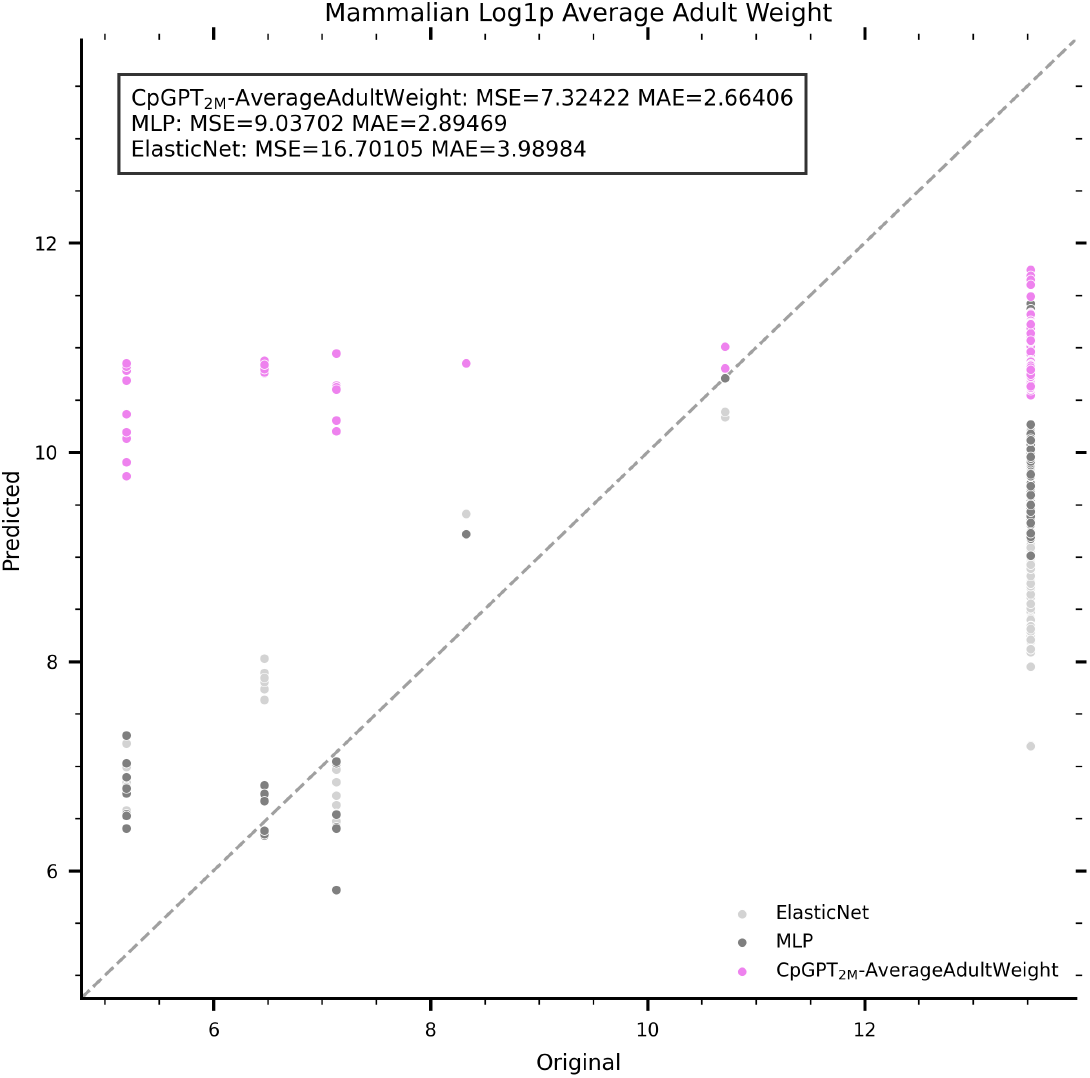
Scatterplot for the prediction of the log1p average adult weight comparing ElasticNet, MLP, and CpGPT_2M_-AverageAdultWeight.

**Supplementary Figure 19:**
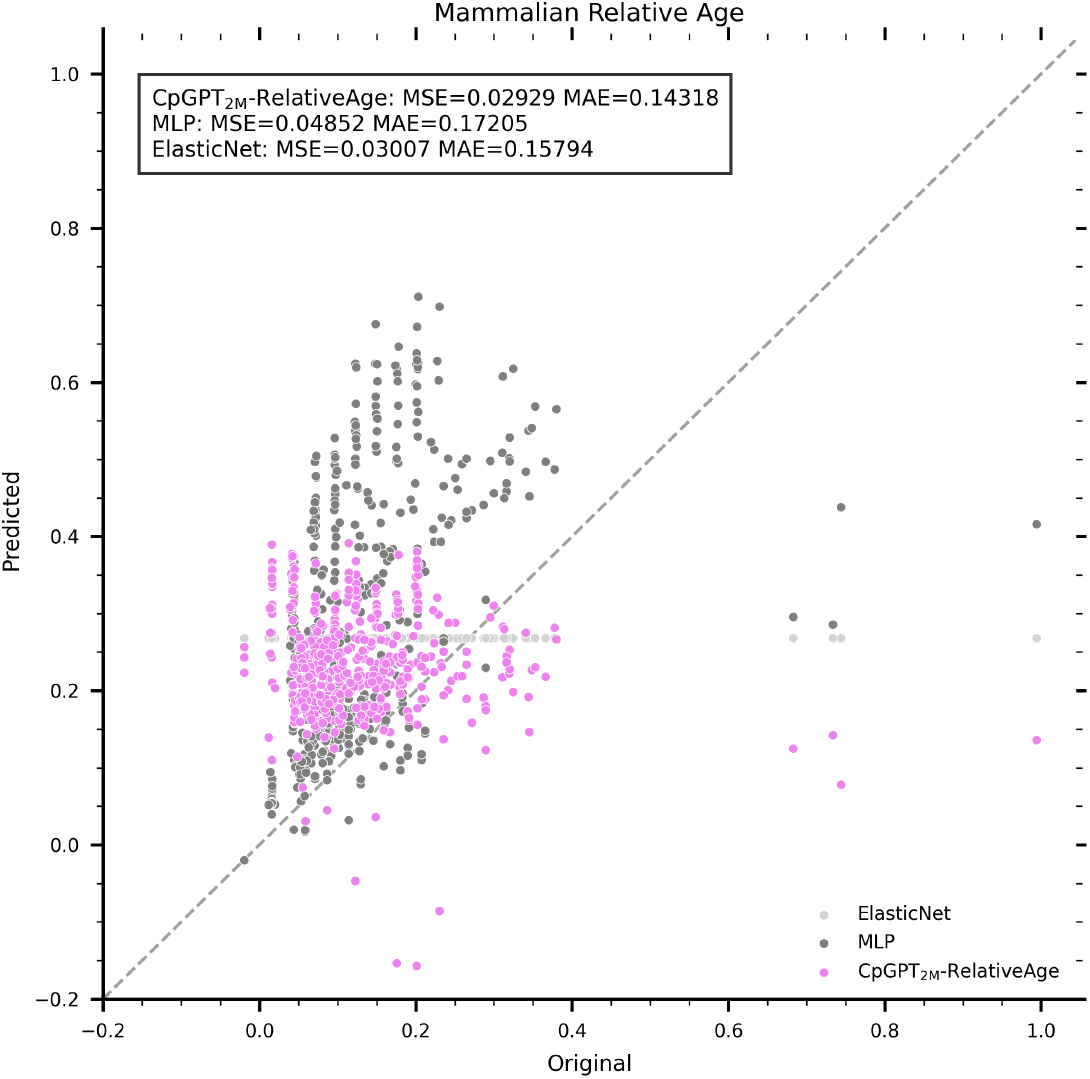
Scatterplot for the prediction of the relative age comparing ElasticNet, MLP, and CpGPT_2M_-RelativeAge.

**Supplementary Figure 20:**
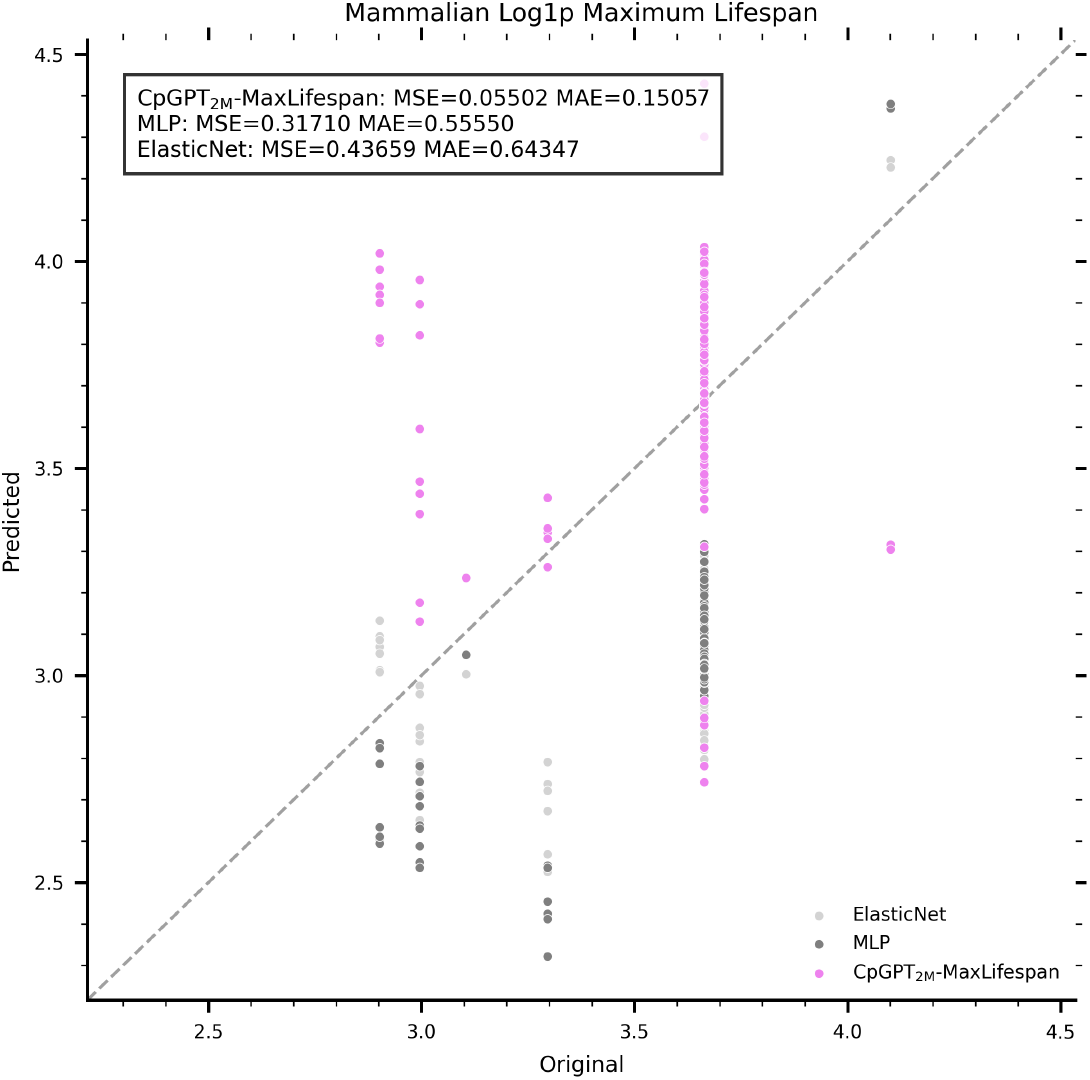
Scatterplot for the prediction of the log1p maximum lifespan comparing ElasticNet, MLP, and CpGPT_2M_-MaxLifespan.

**Supplementary Figure 21:**
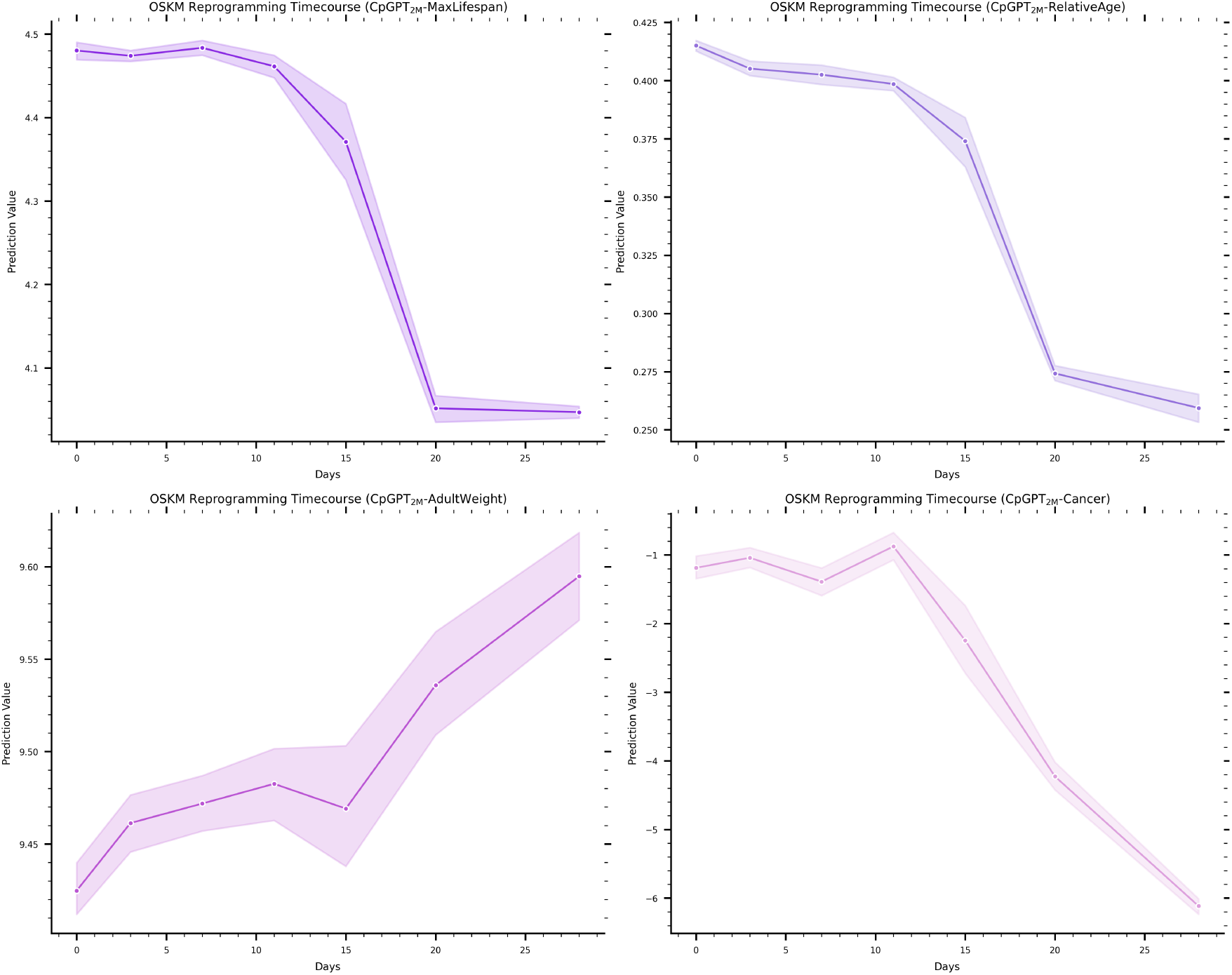
Lineplots showing the 95% confidence interval of five CpGPT models in the OSKM dataset with 10 different random input subsets of CpGs.

**Supplementary Figure 22:**
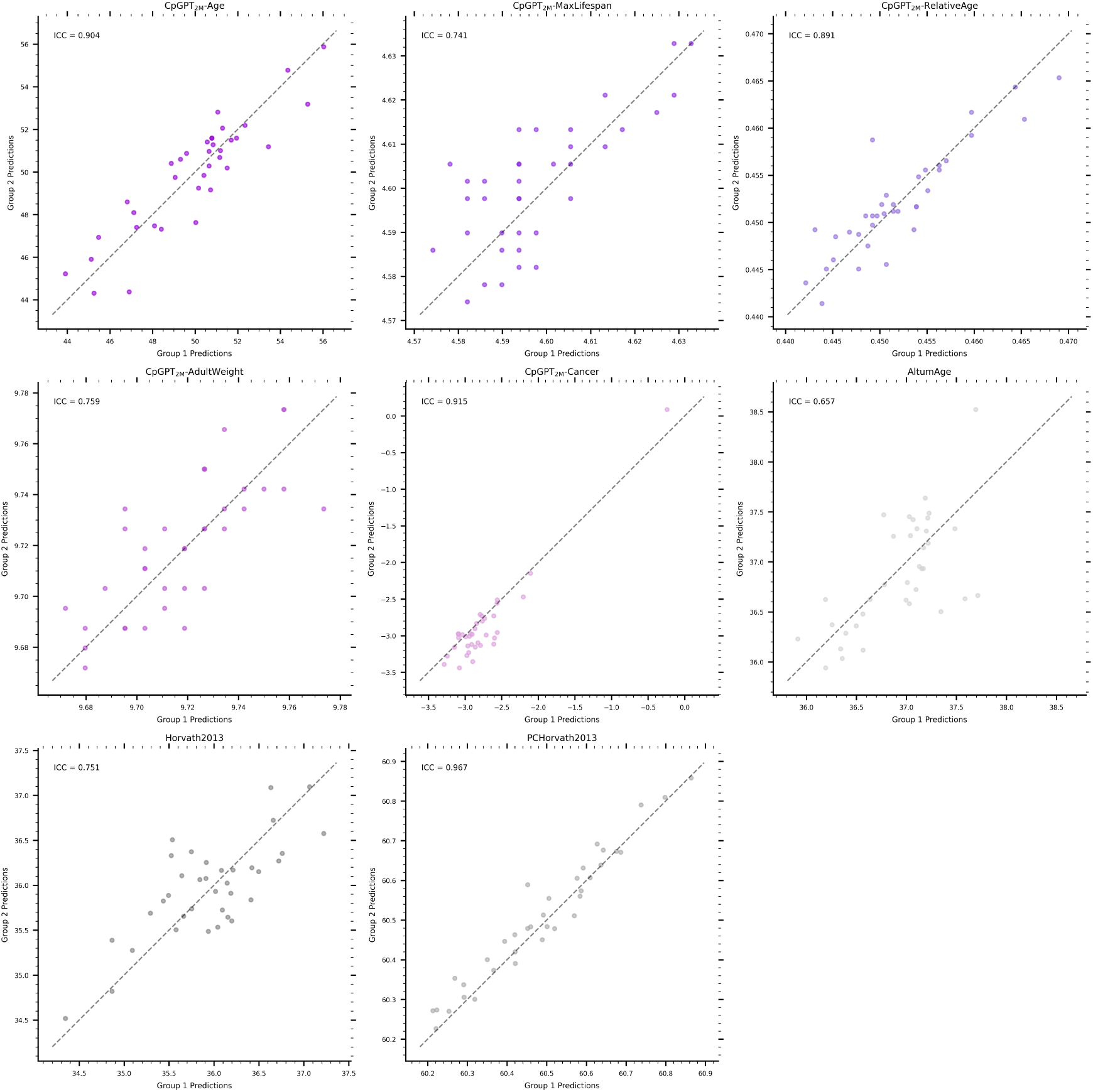
Scatterplot showing the predictions for each technical replicate in the reliability dataset with different CpGPT models and three epigenetic clocks.

## Notes

### Summary of Updates

The affiliations of the authors have now been updated.

